# EFFECTS OF SPINAL TRANSECTION AND LOCOMOTOR SPEED ON MUSCLE SYNERGIES OF THE CAT HINDLIMB

**DOI:** 10.1101/2024.09.19.613891

**Authors:** Alexander N. Klishko, Jonathan Harnie, Claire E. Hanson, S. Mohammadali Rahmati, Ilya A. Rybak, Alain Frigon, Boris I. Prilutsky

## Abstract

It was suggested that during locomotion, the nervous system controls movement by activating groups of muscles, or muscle synergies. Analysis of muscle synergies can reveal the organization of spinal locomotor networks and how it depends on the state of the nervous system, such as before and after spinal cord injury, and on different locomotor conditions, including a change in speed. The goal of this study was to investigate the effects of spinal transection and locomotor speed on hindlimb muscle synergies and their time-dependent activity patterns in adult cats. EMG activities of 15 hindlimb muscles were recorded in 9 adult cats of either sex during tied-belt treadmill locomotion at speeds of 0.4, 0.7, and 1.0 m/s before and after recovery from a low thoracic spinal transection. We determined EMG burst groups using cluster analysis of EMG burst onset and offset times and muscle synergies using non-negative matrix factorization. We found five major EMG burst groups and five muscle synergies in each of six experimental conditions (2 states x 3 speeds). In each case, the synergies accounted for at least 90% of muscle EMG variance. Both spinal transection and locomotion speed modified subgroups of EMG burst groups and the composition and activation patterns of selected synergies. However, these changes did not modify the general organization of muscle synergies. Based on the obtained results, we propose an organization for a pattern formation network of a two-level central pattern generator that can be tested in neuromechanical simulations of spinal circuits controlling cat locomotion.

## Introduction

A common morphological feature of limb design in mammals is muscle redundancy (or abundance), defined as a seemingly excessive number of muscles with respect to the number of kinematic degrees of freedom of the limb. For example, the hindlimbs of rodents, cats and dogs have over 30 muscles serving 7 major degrees of freedom, with 3 at the hip, 1 or 2 at the knee and 2 or 3 at the ankle (Burkholder and Nichols, 2004; Charles et al., 2016; Ramalingasetty et al., 2021; Stark et al., 2021). It has been suggested that the nervous system simplifies its control of multiple muscles by combining them in muscle groups, termed motor modules or synergies, and activating each of these groups with a specific time-dependent activation pattern (Bernstein, 1967; Ivanenko et al., 2005; d’Avella et al., 2006; Bizzi et al., 2008; Lacquaniti et al., 2012; Giszter, 2015; Bernstein, 2021; Cheung and Seki, 2021). Besides synergies revealed at the level of individual muscles, there is ample evidence for the existence of motoneuronal synergies within and across muscles that receive shared or unique synaptic input (De Luca and Erim, 1994; Gibbs et al., 1995; Madarshahian et al., 2021; Del Vecchio et al., 2023; Levine et al., 2023).

Analyses of muscle and motoneuronal synergies during locomotion can potentially reveal the organization of spinal premotor interneuronal networks controlling motoneuronal locomotor activity, called central pattern generators (CPG). Currently, there is little consensus on the organization of mammalian locomotor CPGs with several possible schemes proposed. These include unit burst generators (Grillner, 1981; Grillner and Kozlov, 2021), a two-level CPG comprised of a rhythm generator and a pattern formation network (Burke et al., 2001; McCrea and Rybak, 2008), a nested CPG organization (Berkowitz, 2019), multistable half-center oscillators (Bondy et al., 2016; Parker et al., 2021), or networks with recurrent excitatory and inhibitory connectivity generating rotational dynamics (Linden et al., 2022). Muscle synergies identified in locomotion of rodents (DiGiovanna et al., 2016; Santuz et al., 2019), cats (Krouchev et al., 2006; Markin et al., 2012; Desrochers et al., 2019; Harnie et al., 2021; Klishko et al., 2021), and humans (Ivanenko et al., 2005; Monaco et al., 2010; Saito et al., 2018) indicate the existence of a relatively small number of muscle groups activated together by common inputs, consistent with several versions of CPG organization.

Several factors, such as locomotion speed and the state of the nervous system (e.g. following spinal cord injury), have been reported to affect the number and composition of muscle locomotor synergies. In healthy children (Rozumalski et al., 2017) and younger and older adults (Ivanenko et al., 2003; Monaco et al., 2010; Saito et al., 2018; Dewolf et al., 2019), walking speeds do not affect the number of muscle synergies and have low to moderate effects on the composition and activation patterns of synergies within an age group.

Spinal cord injury appears to affect the number and composition as well as the activation patterns of muscle synergies. For instance, children and adults with incomplete spinal cord injury on average have fewer synergies (2-4) and different synergy composition and activation patterns compared to 4-5 muscle synergies of healthy age-matched humans (Fox et al., 2013; Hayes et al., 2014; Sun et al., 2022). Another study reported 5 muscle synergies in individuals with incomplete spinal cord injury that were similar in composition and patterns to the synergies in a healthy control group (Ivanenko et al., 2003). All the above studies reported greater variability of synergy muscle synergies and their activation patterns in studies of patients with spinal cord injury could be explained by differences in the severity and location of the spinal injury, the number of studied muscles, the type of walking assistance, and rehabilitation interventions. These differences make it difficult to reveal consistent muscle synergies and a potential organization of spinal locomotor CPGs that are likely to operate not only in quadrupedal mammals (McCrea and Rybak, 2008; Kiehn, 2016; Grillner and Kozlov, 2021) but also in humans (Duysens and Van de Crommert, 1998; Shapkova, 2004; Minassian et al., 2023).

Two recent studies in cats with low-thoracic spinal transection (i.e. spinal cats) have revealed up to 7 muscle groups that are activated together in the walking cycle and are generally similar to the groups during intact locomotion (Desrochers et al., 2019; Higgin et al., 2020). These muscle groups were identified based on the burst onset and offset times of the electromyographic (**EMG**) activity of several hindlimb muscles using cluster analysis and provided initial information about the potential organization of spinal locomotor networks. Specifically, in both spinal and intact cats, there are muscle activity burst groups comprised of either flexor or extensor muscles and some of these groups contain muscles operating at different joints (Krouchev et al., 2006; Markin et al., 2012; Desrochers et al., 2019; Higgin et al., 2020; Harnie et al., 2021; Klishko et al., 2021). These results suggest that all muscles within individual flexor and extensor burst groups receive a common input from a flexor and an extensor CPG half-center, respectively. However, the above cluster analysis cannot determine the activity contribution of individual muscles within a group or the pattern of the activation input to each group.

Therefore, the goal of this study was to determine hindlimb muscle synergies and their activation patterns in cats before and after a low-thoracic spinal transection during locomotion at different speeds on a tied-belt treadmill. We tested the hypothesis that a spinal mechanism controls the number, composition and activity patterns of muscle synergies as a function of locomotor speed.

## Methods

### Animals and ethical information

All procedures were approved by the Animal Care Committee of the Université de Sherbrooke (Protocol 442-18) in accordance with policies and directives of the Canadian Council on Animal Care. We obtained the current data set from nine adult purpose-bred cats (> 1 year of age at the time of experimentation), 4 females and 5 males, weighing between 3.4 kg and 4.8 kg, purchased from Marshall BioResources (North Rose, NY, USA). Before and after experiments, cats were housed and fed (weight-dependent metabolic diet and water ad libitum) in a dedicated room within the animal care facility of the Faculty of Medicine and Health Sciences at the Université de Sherbrooke. We followed the ARRIVE guidelines 2.0 for animal studies (Percie du Sert et al., 2020). The investigators understand the ethical principles under which the journal operates, and our work complies with this animal ethics checklist. In order to maximize the scientific output of each animal, they were used in other studies to investigate different scientific questions, some of which have been published (Harnie et al., 2021; Audet et al., 2022; Mari et al., 2023; Harnie et al., 2024).

### Surgical procedures

The implantation and spinal transection surgeries were performed under aseptic conditions with sterilized equipment in an operating room. Prior to surgery, cats were sedated with an intramuscular (i.m.) injection of butorphanol (0.4 mg/kg), acepromazine (0.1 mg/kg), and glycopyrrolate (0.01 mg/kg). We then injected a mixture (0.05 ml/kg, i.m.) of diazepam (0.25 mg/kg) and ketamine (2.0 mg/kg) in a 1:1 ratio five minutes later for induction. We shaved the animal’s fur (back, stomach, fore- and hindlimbs) and cleaned the skin with chlorhexidine soap. Cats were anesthetized with isoflurane (1.5-3%) and O_2_ delivered with a mask and then with a flexible endotracheal tube. The depth of anesthesia was confirmed by applying pressure to a paw (to detect limb withdrawal) and by assessing the size and reactivity of pupils. Isoflurane concentration was adjusted throughout the surgery by monitoring cardiac and respiratory rates. Body temperature was maintained constant (37 ± 0.5°C) using a water-filled heating pad placed under the animal, an infrared lamp placed ∼50 cm over it and a continuous infusion of lactated Ringer’s solution (3 ml/kg/h) through a catheter placed in a cephalic vein. At the end of surgery, we injected subcutaneously with an antibiotic (cefovecin, 8 mg/kg) and a fast-acting analgesic (buprenorphine, 0.01 mg/kg). We also taped a fentanyl (25 µg/h) patch to the back of the animal 2-3 cm rostral to the base of the tail for prolonged analgesia, which we removed 4-5 days later. After surgery, the cats were placed in an incubator and closely monitored until they regained consciousness. We administered another dose of buprenorphine ∼7 hours after surgery.

To record EMG, we directed pairs of Teflon-insulated multistrain fine wires (AS633; Cooner Wire Co., Chatsworth, CA, USA) subcutaneously from two head-mounted 34-pin connectors (Omnetics Connector Corp., Minneapolis, MN, USA). Two wires, stripped of 1–2 mm of insulation, were sewn into the belly of selected hindlimb muscles for bipolar recordings. The head-mounted connectors were fixed to the skull using dental acrylic and four to six screws. We verified electrode placement during surgery by stimulating each muscle through the appropriate head connector channel to assess the biomechanically desired muscle contraction. During experiments, EMG signals were pre-amplified (×10, custom-made system), bandpass filtered (30–1,000 Hz) and amplified (100–5,000×) using a 16-channel amplifier (model 3500; AM Systems, Sequim, WA, USA). EMG data were digitized (5,000 Hz) with a National Instruments (Austin, TX, USA) card (NI 6032E), acquired with custom-made acquisition software and stored on computer. We implanted the following hindlimb muscles bilaterally: tibialis anterior (TA, ankle dorsiflexor), peroneus longus (PLO, ankle abductor and dorsiflexor), flexor digitorum longus (FDL, digits and ankle plantarflexor), soleus (SO, ankle plantarflexor), medial gastrocnemius (MG, ankle plantarflexor and knee flexor), plantaris (PL, digits and ankle plantarflexor and knee flexor), vastus lateralis (VL, knee extensor), sartorius anterior (SRTa, hip flexor and knee extensor), biceps femoris posterior (BFP, hip extensor and knee flexor), semitendinosus (ST, hip extensor and knee flexor), biceps femoris anterior (BFA, hip extensor), gluteus (GLU, hip abductor and extensor), caudofemoralis (CF, hip abductor and extensor), and iliopsoas (IP, hip flexor).

One to two weeks after electrode implantation, we collected data in the intact state for 4-6 weeks. A complete spinal transection was then made at low thoracic levels. Before surgery, we sedated the cat with an intramuscular injection of a cocktail containing butorphanol (0.4 mg/kg), acepromazine (0.1 mg/kg) and glycopyrrolate (0.01 mg/kg) and inducted with another intramuscular injection (0.05 ml/kg) of ketamine (2.0 mg/kg) and diazepam (0.25 mg/kg) in a 1:1 ratio. We shaved the fur overlying the back, stomach, and hindlimbs and cleaned the skin with chlorhexidine soap. The cat was then anesthetized with isoflurane (1.5-3%) and O_2_ using a mask concentration was confirmed and adjusted throughout the surgery by monitoring cardiac and respiratory rates, by applying pressure to the paw to detect limb withdrawal and by assessing muscle tone. Once the animal was deeply anesthetized, the skin was incised over the 12^th^ and 13^th^ (T12-T13) thoracic vertebrae and after setting aside muscle and connective tissue, a small laminectomy of the dorsal bone was made. After exposing the spinal cord, we applied xylocaine (lidocaine hydrochloride, 2%) topically and made two to three injections within the spinal cord. We then completely transected the spinal cord with surgical scissors. We then cleaned the ∼0.5 cm gap between the two cut ends of the spinal cord and stopped any residual bleeding. We verified that no spinal cord tissue remained connecting rostral and caudal ends, which we later confirmed histologically. A hemostatic material (Spongostan) was inserted within the gap, and muscles and skin were sewn back to close the opening in anatomic layers. At the end of surgery, we injected an antibiotic (Cefovecin, 0.1 ml/kg) subcutaneously and taped a transdermal fentanyl patch (25 mcg/h) to the back of the animal 2–3 cm rostral to the base of the tail to provide prolonged analgesia, which was removed 4-5 days later. We also injected buprenorphine (0.01 mg/kg), a fast-acting analgesic, subcutaneously at the end of the surgery and a second dose ∼7 h later. After surgery, we placed the cat in an incubator until it regained consciousness. After spinal transection, we manually expressed the cat’s bladder and large intestine 2-3 times daily. Cats were then monitored daily by experienced personnel. The hindlimbs were cleaned as needed to prevent infection. Following surgery, once the animal was fully conscious (capable of holding its head upright, pupils no longer dilated, calm behaviour), we mixed a can of moist food with water and left it for the animal’s consumption and rehydration. Dry food was then offered the next day. In the 8 hours after waking up, the animal was closely monitored, repositioned and stimulated to eat and drink. To monitor the state of hydration of the animal, we performed skin fold tests, tested for capillary refill time and verified the urine (e.g. colour, consistency) when manually voiding the bladder. If the animal was not sufficiently hydrated, we performed and maintained fluid therapy (200 ml/day of NaCl 0.9%, sub-cutaneous) until hydration returned to normal. The animals were closely monitored, and their general state (respiratory and cardiac rates, temperature and overall behaviour) was evaluated by qualified personnel three times per day. At 5-7 days post-transection, animals were fully independent for feeding, drinking and maintaining their body temperature within normal values (37°C ± 0.5), as well as moving around using their forelimbs. At the end of experiments, cats were anaesthetized with isoflurane (1.5–3.0%) and O2 before receiving a lethal dose of pentobarbital (120 mg/kg) through the left or right cephalic vein. Cardiac arrest was confirmed using a stethoscope to determine the death of the animal.

### Experimental design and data collection

Data collection in spinal cats began between the 8th and 15th weeks after spinal transection, based on the recovery of a robust hindlimb locomotion in the forward direction with full hindquarter weight support without or with perineal stimulation. For perineal stimulation, the experimenter manually pinched or rubbed the skin under the tail with the index finger and thumb. During data collection in spinal cats, the experimenter held the tail of the animal to provide mediolateral balance assistance but did not provide weight support. We did not train spinal cats to recover hindlimb locomotion as discussed previously (Harnie et al., 2019). Cats performed consisting of two independently controlled running surfaces 120 cm long and 30 cm wide (Bertec, Columbus, OH, USA). A Plexiglas separator (120 cm long, 3 cm high, and 0.5 cm wide) was placed between the left and right belts to prevent the limbs from impeding each other. Cats performed tied-belt locomotion (equal left-right speeds) at speeds of 0.1 m/s to 1.0 m/s in 0.1 m/s increments. At each speed, the objective was to obtain a minimum of 10-15 consecutive cycles.

We collected kinematic data by capturing videos of the left and right sides using two cameras (Basler AcA640-100 g, Basler AG, Germany) at 60 frames/s with a spatial resolution of 640 x 480 pixels. A custom-made program (LabVIEW, National Instruments, Austin, TX, USA) acquired the images and synchronized acquisition with EMG data. EMG signals were pre-amplified (10×, custom-made system), bandpass filtered (30–1000 Hz), and amplified (100–5000×) using a 16- channel amplifier (model 3500; A-M Systems). As we implanted more than 16 muscles per cat, we obtained data in each locomotor condition twice, one for each connector, as our data acquisition system is limited to 16 channels. EMG data were digitized (2000 Hz) with a National Instruments card (NI 6032E, Austin, TX, USA), acquired with custom-made acquisition software and stored on computer.

### Analysis of muscle activity, burst groups and synergies

We described the analysis of EMG activity and determination of EMG burst groups and muscle synergies previously (Harnie et al., 2021; Klishko et al., 2021) and therefore provide only a short summary here. We rectified raw EMG signals and determined onset and offset times of each burst using an EMG threshold (typically the mean EMG interburst baseline plus 2 standard deviations, SD). We normalized EMG burst onset and offset times to the duration of the corresponding cycle duration and performed a cluster analysis using the minimum spanning tree algorithm in MATLAB (MathWorks, Natick, MA, USA) for each combination of spinal cord condition (intact and spinal-transected) and locomotion speed (0.4, 07, and 1.0 m/s). We computed the mean amplitude of each EMG burst and normalized it to the maximum mean burst magnitude determined for each EMG channel and cat across all experimental conditions.

To determine muscle synergies, we first subtracted EMG values below the burst threshold from the raw rectified EMG within each trial and smoothed the signal using a Butterworth 4^th^ order zero-lag digital filter with a cutoff frequency of 10 Hz. We used a negative matrix factorization algorithm (NNMF; MATLAB function *nnmf* with parameter *replicates* = 20) to compute muscle weights (the relative contribution of each muscle to individual synergies, matrix **W**) and time-dependent activation of each synergy (matrix **C**) from the following equation: **D** = **WC** + **error**. Here **D** is a (*m* × *T*) matrix of EMG envelope values for each muscle and percentage of locomotion cycle, *m* = 15 is the number of muscles, *T* = 100 is the number of normalized time samples in the cycle; **W** is a (*m* × *n*) matrix of muscle weights indicating each muscle contribution to each synergy, *n* is the number of synergies; **C** is a (*n* × *T*) matrix of time-dependent activation coefficients for each synergy; **error** is a (*m* × *T*) matrix of deviations of matrix **D** from matrix **D\*** = **WC**. We generated 50 matrices **D** by randomly selecting cycles of EMG envelopes for each muscle across all cats with available cycles within each experimental condition (Table 1) to maximize representation of different cats in computed synergies (Klishko et al., 2021). We computed matrices **W** and **C** for each experimental condition using a different number of synergies accounts for at least 90% of the mean EMG variance computed across all 50 matrices **D** within each condition.

**Table 1.** Animal characteristics and number of investigated locomotion cycles in all experimental conditions.

| Cat | Sex | Mass,<br>kg | Experimental conditions |  |  |  |  |  | Total |
| --- | --- | --- | --- | --- | --- | --- | --- | --- | --- |
|  |  |  | Intact<br>0.4 m/s | Intact<br>0.7 m/s | Intact<br>1.0 m/s | Spinal<br>0.4 m/s | Spinal<br>0.7 m/s | Spinal<br>1.0 m/s |  |
| AM | F | 3.6 | 20 | 42 | 29 | 60 | 51 | 33 | 235 |
| C3 | M | 4.7 | 45 | 59 | 46 | 29 | 62 | 28 | 269 |
| CR | F | 4.1 | 28 | 31 | 27 | 12 | 15 | 16 | 129 |
| DA | M | 4.4 | 39 | 53 | 58 | 44 | 39 | 38 | 271 |
| DO | F | 3.6 | 50 | 71 | 76 | 60 | 69 | 75 | 401 |
| HA | M | 4.0 | 51 | - | - | 51 | 34 | 40 | 176 |
| SA | F | 3.6 | 46 | 57 | - | 132 | 73 | 76 | 384 |
| SN | M | 4.8 | 64 | 79 | 72 | 59 | 60 | 40 | 374 |
| TO | M | 3.4 | 31 | 36 | 30 | 27 | 65 | 29 | 218 |
| Mean<br>±SD or<br>Total |  | 4.02<br>±<br>0.51 | 374 | 428 | 338 | 474 | 468 | 375 | 2457 |

To evaluate the similarity of muscle synergies in different experimental conditions, we also computed matrices **W**_COM_ and **C**_COM_ for different combinations of spinal states and speeds and evaluated the variance accounted for by their products 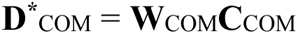 in the corresponding experimental EMG patterns **D**_COM_ (Harnie et al., 2021; Klishko et al., 2021). Here the subscript COM denotes a specific combination of spinal states and speeds. For example, we evaluated the effect of spinal cord state on muscle synergies at given locomotion speeds by computing matrices 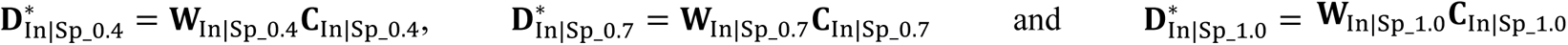 and calculated the variance accounted for by these matrices 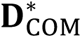, containing reconstructed EMG patterns from muscle synergies, with the corresponding experimental EMG patterns (matrices 𝐃_In_0.4_, 𝐃_Sp_0.4_, etc.). The subscript In|Sp in the above example indicates a combination of the intact and spinal states; the subscripts 0.4, 0.7, and 1.0 correspond to locomotion speeds. In other words, matrices 𝐃_In|Sp_0.4_, 𝐃_In|Sp_0.7_, and 𝐃_In|Sp_1.0_ are composed of all intact and spinal EMG patterns for each speed (Fig. 1 top, middle, and bottom panels, respectively). The dimensions of these and corresponding **C**_COM_ matrices are (*m* × 2*T*). The dimensions of the corresponding matrices **W**_COM_ are (*m* × *n*). Similarly, we evaluated the effect of locomotion speed on muscle synergies in the intact and spinal states by computing matrices 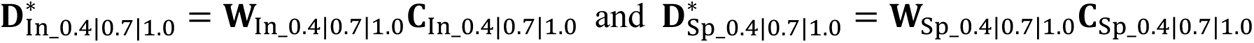, where the subscript 0.4|0.7|1.0 indicates a combination of all locomotor speeds together. The corresponding matrices of experimental EMG patterns (𝐃_In_0.4|0.7|1.0_ and 𝐃_Sp_0.4|0.7|1.0_) are composed of all EMG patterns on the left and right panels of Fig. 1, respectively. The dimensions of these and the corresponding **C**_COM_ matrices are (*n* × 3*T*); the corresponding matrices 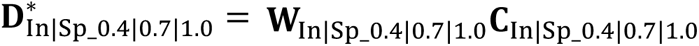 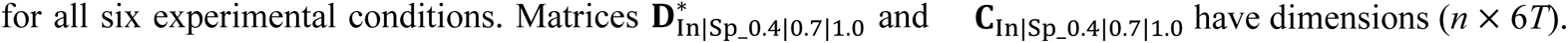.

**Figure 1.**
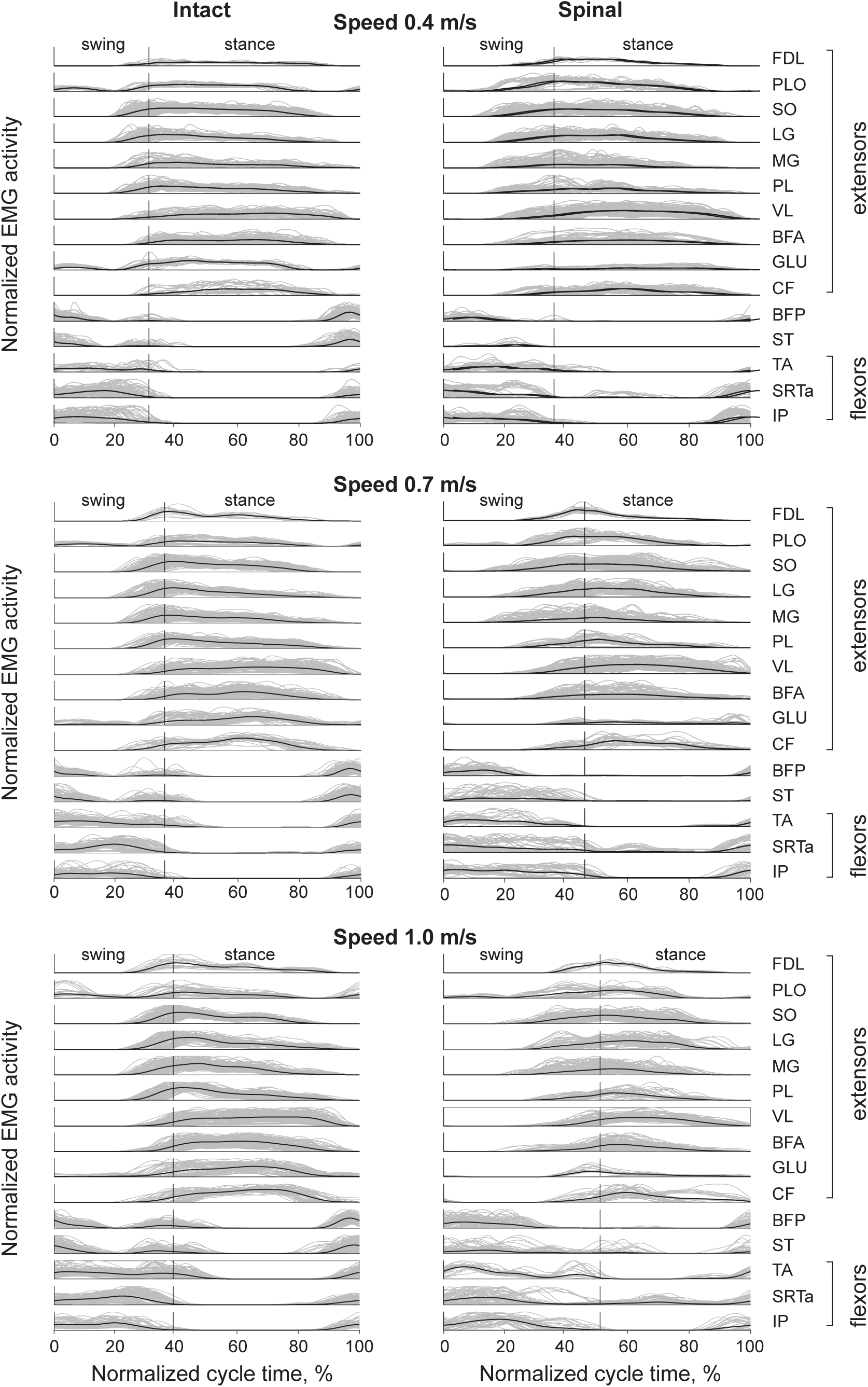
Low-pass filtered patterns of EMG activity of hindlimb muscles during intact (left panels) and spinal (right panels) tied-belt treadmill locomotion at speeds 0.4 m/s (top panels), 0.7 m/s (middle panels), and 1.0 m/s (bottom panels). Gray lines show EMG patterns of all analyzed 2457 cycles of 9 cats (see Table 1), thick black lines are the mean EMG patterns across all cycles and cats within a muscle and experimental condition. The EMG magnitude of each muscle is normalized to the peak low-pass filtered EMG across all experimental conditions within the cat. The vertical axis in each plot designates the range of the normalized EMG magnitude from 0 to 1. The vertical line within each EMG panel separates the swing and stance phases. Extensors are muscles with a stance phase related activity and primary function of supporting body against gravity and propelling it forward: FDL, flexor digitorum longus (digits and ankle plantarflexor); PLO, peroneus longus (ankle abductor and a dorsiflexor); SO, soleus (ankle plantarflexor); MG, medial gastrocnemius (ankle plantarflexor and knee flexor); PL, plantaris (digits and ankle plantarflexor and knee flexor); VL, vastus lateralis (knee extensor); BFA, biceps femoris (hip extensor); GLU, gluteus (hip abductor and extensor); CF, caudofemoralis (hip abductor and extensor). Flexors are muscles with a swing phase related activity: TA, tibialis anterior (ankle dorsiflexor); SRTa, sartorius anterior (hip flexor and knee extensor); IP, iliopsoas (hip flexor). Two muscles (BFP, biceps femoris posterior, hip extensor and knee flexor) and ST (semitendinosus, hip extensor and knee flexor) are bifunctional muscles with swing and stance related activity.

We also compared similarities of muscle contributions to each synergy between different states (intact vs spinal) and locomotion speeds by computing the scalar product (the angle) between pairs of m-dimensional vectors of muscle weights (rows in **W**) for different combinations of experimental conditions (Cheung et al., 2005; Klishko et al., 2021).

### Statistics

We evaluated the significance of the effects of independent factors of state (intact, spinal), locomotion speed (0.4, 0.7, 1.0 m/s), and/or muscle (*m* = 15) on dependent variables, such as temporal cycle characteristics, normalized EMG burst magnitude, and onset and offset times. In this statistical analysis, we considered animals as a random factor. We evaluated significance of main effects of the above factors and their interactions using a linear mixed-effects model (MIXED function; IBM SPSS Statistics 29, Chicago, IL, USA). We performed post hoc pairwise comparisons with Bonferroni adjustments.

We used a similar linear mixed-effects model to test for the significance of the effects of state and locomotion speed on muscle weights (matrices **W**) and the angle between pairs of muscle weight vectors for different combinations of experimental conditions within synergies. A separate linear mixed-effects model analysis was performed to evaluate significance of the difference in synergy activation coefficients (matrices **C**) at each 10%-time bin of the locomotor cycle. In addition, we computed the coefficient of determination R^2^ between pairs of the mean activation patterns corresponding to each experimental condition (e.g., 𝐂_In_0.4_, 𝐂_Sp_1.0_, etc.) and the corresponding components of matrices **C**_COM_ (e.g., 𝐂_In_0.4|0.7|1.0_, 𝐂_InSp_1.0_, etc.). For all statistical tests, the significance was set at 0.05.

## Results

### General characteristics of EMG activity patterns and bursts

Figure 1 demonstrates experimentally obtained EMG linear envelopes of 15 hindlimb muscles in analyzed locomotor cycles in the intact and spinal states at treadmill speeds of 0.4, 0.7 and 1.0 m/s. Hindlimb muscles with extensor (stance phase) related activity (top 10 muscles in each panel) initiated their activity at 10-15% and 20-25% of the cycle time prior to stance onset in intact and spinal conditions, respectively. The extensor activity bursts ended in the last 10-20% of the stance phase. Hindlimb muscles with flexor (swing phase) related activity (bottom three muscles) initiated their activity 10-20% prior to swing onset and the activity typically lasted until the end of swing with little or no-coactivation with extensor muscles. Bifunctional BFP and ST muscles had flexor and extensor bursts at the stance-swing and swing-stance transitions, respectively, during intact locomotion at all speeds. EMG patterns of these muscles changed in the spinal state as the extensor burst became much smaller or disappeared, while the flexor burst increased in duration, especially in ST. Two muscles with stance related activity, PLO and GLU, also had a flexor burst PLO is an ankle abductor and dorsiflexor (Young et al., 1993) and GLU is a hip abductor and extensor (Crouch, 1969).

The normalized extensor burst magnitude across all muscles was generally greater in intact locomotion compared to spinal locomotion at 0.4 m/s (mean ± SD: 0.54 ± 0.21 vs 0.50 ± 0.25), 0.7 m/s (0.60 ± 0.21 vs 0.52 ± 0.27), and 1.0 m/s (0.71 ± 0.22 vs 0.55 ± 0.25) (Fig. 2A). The extensor burst magnitude also increased with locomotor speed. The effects of the state, speed, and muscle independent factors were significant (F_1,5475_ = 268.6, p < 0.001, F_2,5475_ = 133.3, p < 0.001, and F1_12,5475_ = 148.3, p < 0.001, respectively). The interaction effect was also significant (F_60,5475_ = 24.8, p < 0.001). We found no significant differences between the intact and spinal states for the extensor bursts of FDL (p = 0.891, p = 0.809), LG (p = 0.783, p = 0.986), and CF (p = 0.313, p = 0.455) at speeds 0.4 and 0.7 m/s (Fig. 2A). In only one case did we observe an extensor muscle with a larger burst in the spinal state, the VL at a speed 0.4 m/s. The extensor bursts of PLO had a greater magnitude in the spinal state compared to the intact state at all speeds (p < 0.001). The SRTa did not have an extensor burst in the intact state at a speed of 0.4 m/s, the SRTa extensor burst magnitude was the same in the intact and spinal state at a speed of 0.7 m/s (p = 0.477), and its magnitude was larger after spinal transection compared to the intact state at 1.0 m/s (p < 0.001). Extensor bursts started before paw contact with the treadmill and terminated before or at the end of the stance phase (Fig. 1). The burst onset times normalized to the cycle times were significantly affected by state (F_1,5479_ = 106.1, p < 0.001), speed (F_2,5479_ = 420.7, p < 0.001), and muscle (F_12,5479_ = 207.4, p < 0.001). For example, the relative burst onset times occurred significantly later in the cycle in the spinal state compared to intact at locomotor speeds of 0.4 m/s (0.29 ± 0.09 vs 0.28 ± 0.06, p < 0.001), 0.7 m/s (0.35 ± 0.10 vs 0.32 ± 0.07, p < 0.001), and 1.0 m/s (0.40 ± 0.10 vs 0.35 ± 0.08, p < 0.001). (Note that the normalized swing duration is substantially longer in the spinal than in intact state (see below), which creates an appearance of an earlier burst onset in the spinal state.) The extensor burst onset times also occurred later in the cycle with increasing speed (F_2,5479_ = 542.8, p < 0.001). The normalized extensor burst offset times significantly depended on the speed, muscle, and condition-speed-muscle interaction (F_2,5466_ = 46.2, p < 0.001; F_12,5468_ = 1073.1, p < 0.001; and F_60,5465_ = 19.6, p < 0.001, respectively) but not on the condition (F_1,5472_ = 1.7, p = 0.187). The normalized offset times of extensor bursts occurred later in the cycle in the spinal state compared to intact at speeds 0.4 m/s and 1.0 m/s (0.76 ± 0.12 vs 0.73 ± 0.12, p < 0.001 and 0.77 ± 0.12 vs 0.75 ± 0.12, p = 008, respectively) but not at 0.7 m/s (0.76 ± 0.12 vs 0.75 ± 0.12, p = 790).

**Figure 2.**
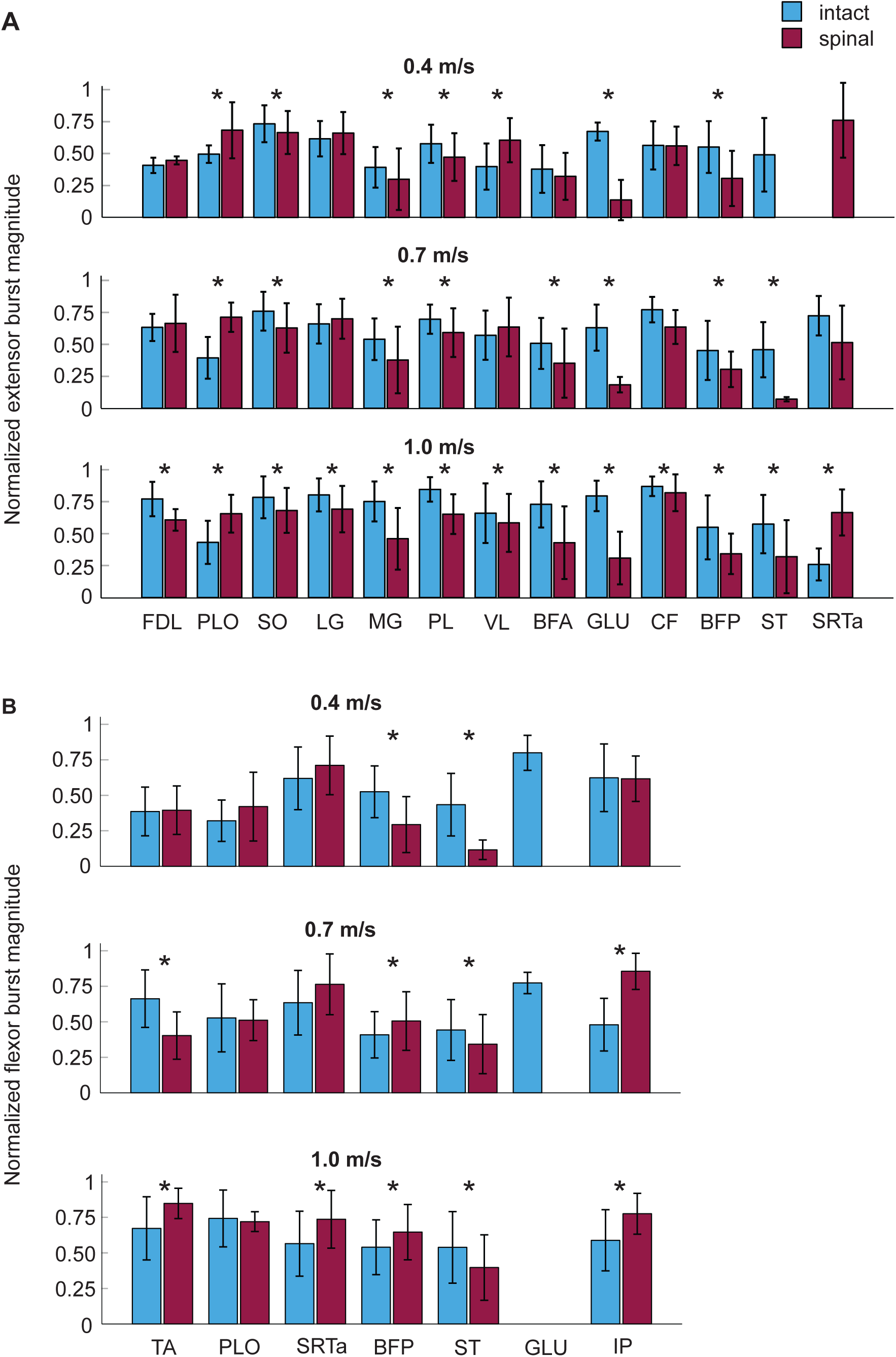
Normalized EMG burst activity (mean ± SD) in intact and spinal states during locomotion at speeds 0.4, 0.7 and 1.0 m/s. **A**: Normalized EMG activity of extensor (stance related) bursts. **B**: Normalized EMG activity of flexor (swing related) bursts. Asterisks indicate statistical significance between intact and spinal conditions. For extensor bursts main effect of condition was significant (p < 0.001, F_1,5475_ = 268.6, linear mixed-effects model). For flexor bursts the main effect of condition was not significant (F_1,2035_ = 2.427, p = 0.119), whereas the interaction effect of condition-speed-muscle was significant (p < 0.001). For muscle abbreviations see the legend for Fig. 1.

The main effect of state on the mean amplitude of flexor bursts was not significant (F_1,2035_ = 2.427, p = 0.119), whereas speed, muscle, and state-speed-muscle interactions significantly affected flexor burst magnitude (F_2,2035_ = 82.1, p < 0.001, F_6,2035_ = 78.6, p < 0.001, and F_28,2035_ = 14.6, p < 0.001, respectively). The majority of significant differences in mean amplitude between states occurred at a speed of 1.0 m/s, as most flexor bursts were greater after spinal transection (p < 0.001), except for (**ST**), which was smaller in the spinal state (p < 0.001), and for PLO whose magnitude was not significantly different (p = 0.709) (Fig. 2B). At slower speeds, fewer differences in burst magnitude between states could be seen and they often had the opposite signs, e.g. TA at speed 0.7 m/s (p < 0.001) and BFP at speed 0.4 m/s (p < 0.001). At the slowest speed of 0.4 m/s, only two flexor bursts differed between states (BFP and ST) and both were greater in the intact state. The GLU flexor burst appeared only in the intact state and at two speeds, 0.4 and 0.7 m/s (Figs. 1 and 2**B**).

The flexor bursts typically started prior to swing onset and terminated at or slightly after swing offset (Fig. 1). The normalized flexor burst onset times dependent on the state, locomotion speed, muscle, and their interaction (F_1,2013_ = 160.9, p < 0.001; F_2,2018_ = 3.4, p = 0.035; F_6,2005_ = 59.3, p < 0.001; and F_28,2017_ = 35.6, p < 0.001; respectively). Flexor bursts started prior to swing onset in the intact state at all speeds (0.4 m/s: -0.07 ± 0.04; 0.7 m/s: -0.60 ± 0.04; and 1.0 m/s: -0.07 ± 0.04), whereas in the spinal state the onset started later and closer in time to swing onset (0.4 m/s: -0.04 ± 0.06; 0.7 m/s: 0.003 ± 0.08; and 1.0 m/s: 0.01 ± 0.08). Flexor burst offset times were significantly affected by speed (F_2,2017_ = 69.4, p < 0.001), muscle (F_6,2010_ = 332.2, p < 0.001), and state-speed- muscle interaction (F_28,2017_ = 61.6, p < 0.001), but not by state (F_1,2017_ = 0.86, p = 0.855). Flexor burst offset times did not depend on state at speeds of 0.4 and 1.0 m/s (0.17 ± 0.09 vs 0.21 ± 0.10, p = 0.539 and 0.23 ± 0.14 vs 0.23 ± 0.12, p = 0.479, in the intact and spinal state, respectively); at 1.0 m/s, the flexor bursts terminated later in the cycle in the spinal than in intact state (0.25 ± 0.13 vs 0.19 ± 0.12, p < 0.001).

Flexor and extensor burst normalized timings were also related to the durations of the stance and swing phases (Fig. 1). Figure 3 shows the durations of the cycle, stance and swing phases as well as the duty cycle for all cats and experimental conditions. The cycle time was shorter in the spinal state compared to intact at all three speeds (0.4 m/s: 0.768 ± 0.130 s vs 1.070 ± 0.144 s, p < 0.001; 0.7 m/s: 0.656 ± 0.132 s vs 0.790 ± 0.070 s, p < 0.001; 1.0 m/s: 0.572 ± 0.079 vs 0.637 ± 0.051, p < 0.001) (Fig. 3A). The effects of state, speed, and their interaction were significant (F_1,2444_ = 1754.3, p < 0.001; F_2,2444_ = 2383.4, p < 0.001; F_2,2444_ = 369.4, p < 0.001; respectively). Because stride length is the product of cycle duration and locomotor speed, the mean stride length was computed for each experimental condition from the above mean cycle durations. Stride length was shorter in the spinal state compared to intact and increased with locomotor speed. It was 0.307 m for spinal vs 0.428 m for intact at 0.4 m/s, 0.459 m vs 0.553 m at 0.7 m/s, and 0.572 m vs 0.637 m at 1.0 m/s, respectively.

**Figure 3.**
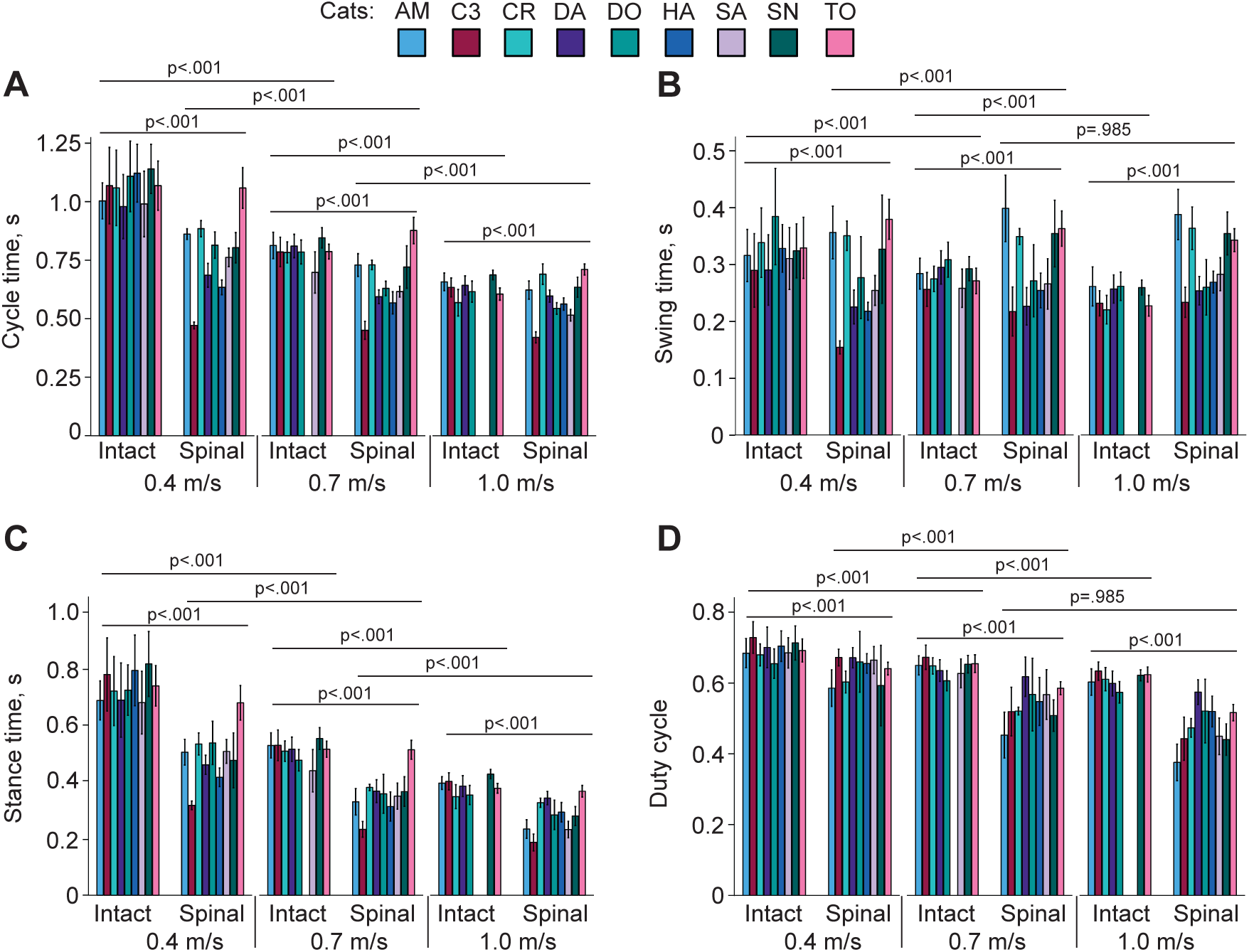
Temporal characteristics of locomotion cycles of individual cats (mean ± SD). **A**: Cycle time. **B**: Swing time. **C**: Stance time. **D**: Duty cycle. Horizontal lines and p-values above pairs of experimental conditions indicate significance of the difference between the conditions.

Swing duration was significantly affected by state and speed and their interaction (F_1,2444_ = 8.4, p = 0.004; F_2,2444_ = 67.2, p < 0.001; and F_2,2444_ = 192.4, p < 0.001; respectively). In the intact state, swing duration decreased with increasing speed from 0.325 ± 0.065 s to 0.283 ± 0.033 s (p < 0.001) and to 0.251 ± 0.027 s (p < 0.001), respectively (Fig. 3B). In the spinal state, swing duration increased from 0.277 ± 0.077 s at 0.4 m/s to 0.299 ± 0.078 s at speed 0.7 m/s (p < 0.001). The swing duration at 1.0 m/s was not significantly different from 0.7 m/s (0.296 ± 0.059 s, p = 0.985). State, speed, and their interaction significantly affected stance duration (F_1,2444_ = 2825.2, p < 0.001; F_2,2444_ = 2915.9, p < 0.001; and F_2,2444_ = 189.9, p < 0.001; respectively), which decreased with increasing speed in both states and was smaller in the spinal state compared to intact (Fig. 3C). Stance durations in the intact and spinal states were 0.745 ± 0.121 s and 0.491 ± 0.090 s at 0.4 m/s, respectively; 0.508 ± 0.059 s and 0.358 ± 0.089 s at 0.7 m/s; and 0.386 ± 0.40 s and 0.276 ± 0.60 s at 1.0 m/s. The duty cycle, the ratio of the stance duration over the cycle duration was significantly affected by state, speed, and their interaction (F_1,2444_ = 1387.7, p < 0.001; F_2,2444_ = 875.0, p < 0.001; and F_2,2444_ = 85.3, p < 0.001; respectively). The duty cycle was significantly smaller in the spinal state compared to intact and decreased with increasing speed in both states respectively at 0.4 m/s; 0.642 ± 0.036 and 0.545 ± 0.079 at 0.7 m/s; and 0.606 ± 0.035 and 0.482 ± 0.078 at 1.0 m/s. A duty cycle below 0.5 indicates a running gait (Hildebrand, 1989). Thus, a walk-to-run transition occurred in spinal cats at a speed between 0.7 m/s and 1.0 m/s, while intact cats performed a walking gait in the entire range of studied speeds up to 1.0 m/s. In the spinal state at 1.0 m/s, the duty cycle was between 0.38 ± 0.05 and 0.47 ± 0.04 in 5 cats (running gait), 0.50 ± 0.09 in one cat (walk-run transition), and between 0.51 ± 0.02 and 0.58 ± 0.03 in 3 cats (walking). The general kinematic characteristics of the locomotor cycle described above are consistent with those described in previous studies in intact and spinal cats (Halbertsma, 1983; Frigon et al., 2014; Frigon et al., 2017), including a decrease in cycle and stance durations with increasing speed and an earlier walk-to-run transition in spinal cats.

### EMG burst groups

We used the minimum spanning tree algorithm in MATLAB to determine EMG burst groups of 15 muscles. Based on the obtained burst clusters and previous similar EMG burst group analyses (Krouchev et al., 2006; Markin et al., 2012; Klishko et al., 2021), we defined 4 or 5 major EMG burst groups and their subgroups in each experimental condition (**Figs. 4-6**). Group 1 consisted of extensor bursts of BFP and ST, except in spinal locomotion at 0.4 m/s and 1.0 m/s where these bursts were combined by the clustering algorithm with the bursts of flexors TA, SRTa, and IP (Figs. 4B, 6B). Group 1 was located at the swing-stance transition on the burst onset-offset time plots (see also Fig. 1). Group 2 consisted of bursts of flexors TA, STRa and IP (subgroup 2_1_) and flexor bursts of GLU_F_ and PLO_F_ (subgroup 2_2_) when present, with all these bursts occurring during swing. During intact locomotion at 1.0 m/s, subgroup 2_1_ was separated into smaller subgroups TA and SRTa_F_-IP (Fig. 6A). All group 2 flexor bursts occurred during swing (Fig. 1). Burst group 3 consisted of extensor bursts of BFP_E_ and ST_E_ (in spinal state at speed 0.4 m/s ST_E_ burst was missing) and occurred at the swing-stance transition. The burst group 4 combined EMG bursts of all hindlimb extensors and extensor bursts of bifunctional PLO and GLU. In some experimental conditions (intact at 0.4 and 1.0 m/s and spinal at 0.7 m/s), bursts of single muscles (BFA and CF, Fig. 4A; FDL, Fig. 5B; PLO and VL, Fig. 6A) were separated into distinct subgroups. In spinal cats at speeds of 0.7 and 1.0 m/s, burst group 4 was combined with extensor group 5 (**Figs. 5B and 6B**). Group 5 consisted of the SRTa extensor burst, which occurred in all experimental conditions except one (intact at 0.4 m/s), but was clustered separately from group 4 in spinal locomotion at 0.4 m/s and intact locomotion at 0.7 and 1.0 m/s. Burst group 5 occurred late in stance and was similar to burst group 5 identified previously during level and upslope overground walking (Klishko et al., 2021) and included rectus femoris (knee extensor, hip flexor).

**Figure 4.**
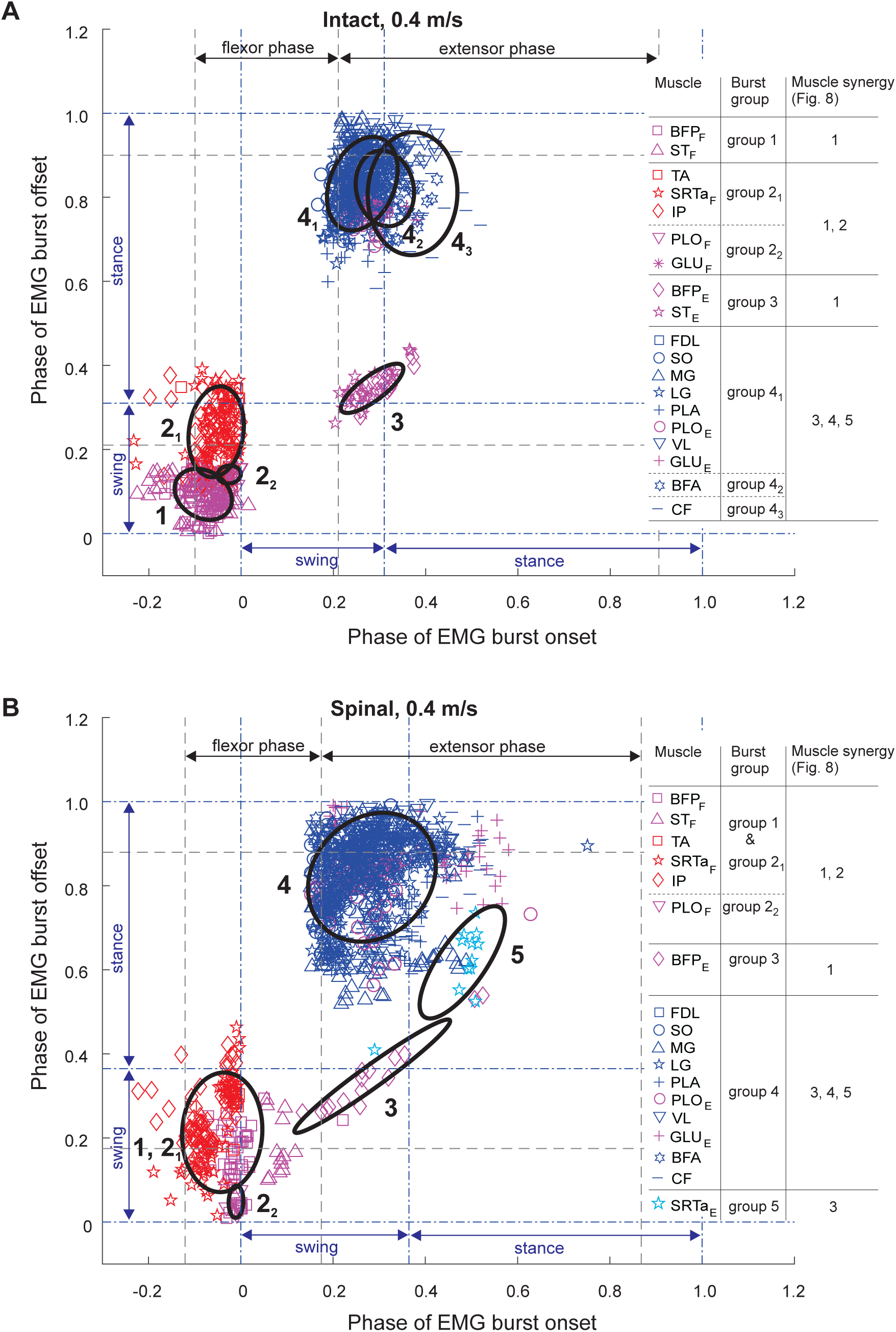
Scatter plots of EMG burst onset and offset times of 15 hindlimb muscles and EMG burst groups for locomotion speed of 0.4 m/s in intact (panel **A**) and spinal (panel **B**) states. Onset and offset times are shown as cycle phase from 0 (swing phase onset) to 1.0 (next swing phase onset). Ellipses and corresponding color-coded symbols denote EMG burst groups/subgroups identified by cluster analysis. The gray dashed lines indicate the onset of the flexor and extensor phases defined by the mean EMG burst onset time minus one SD of IP and SO, respectively. The blue dashed-dotted lines separate the swing and stance phases. For muscle abbreviations see the legend for Fig. 1.

**Figure 5.**
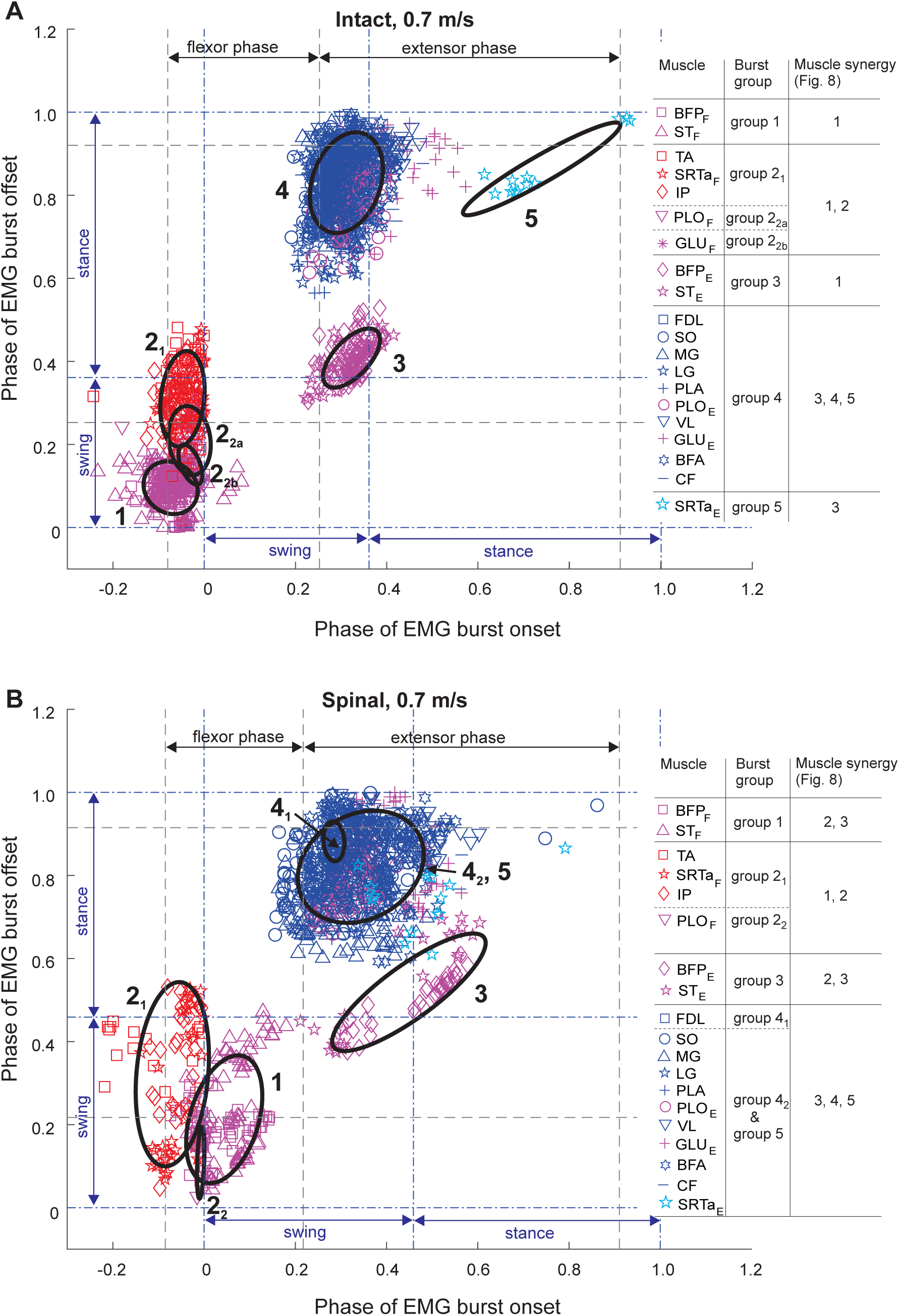
Scatter plots of EMG burst onset and offset times of 15 hindlimb muscles and EMG burst groups for locomotion speed of 0.7 m/s in intact (panel **A**) and spinal (panel **B**) states. For further information see the legend for Fig. 4.

**Figure 6.**
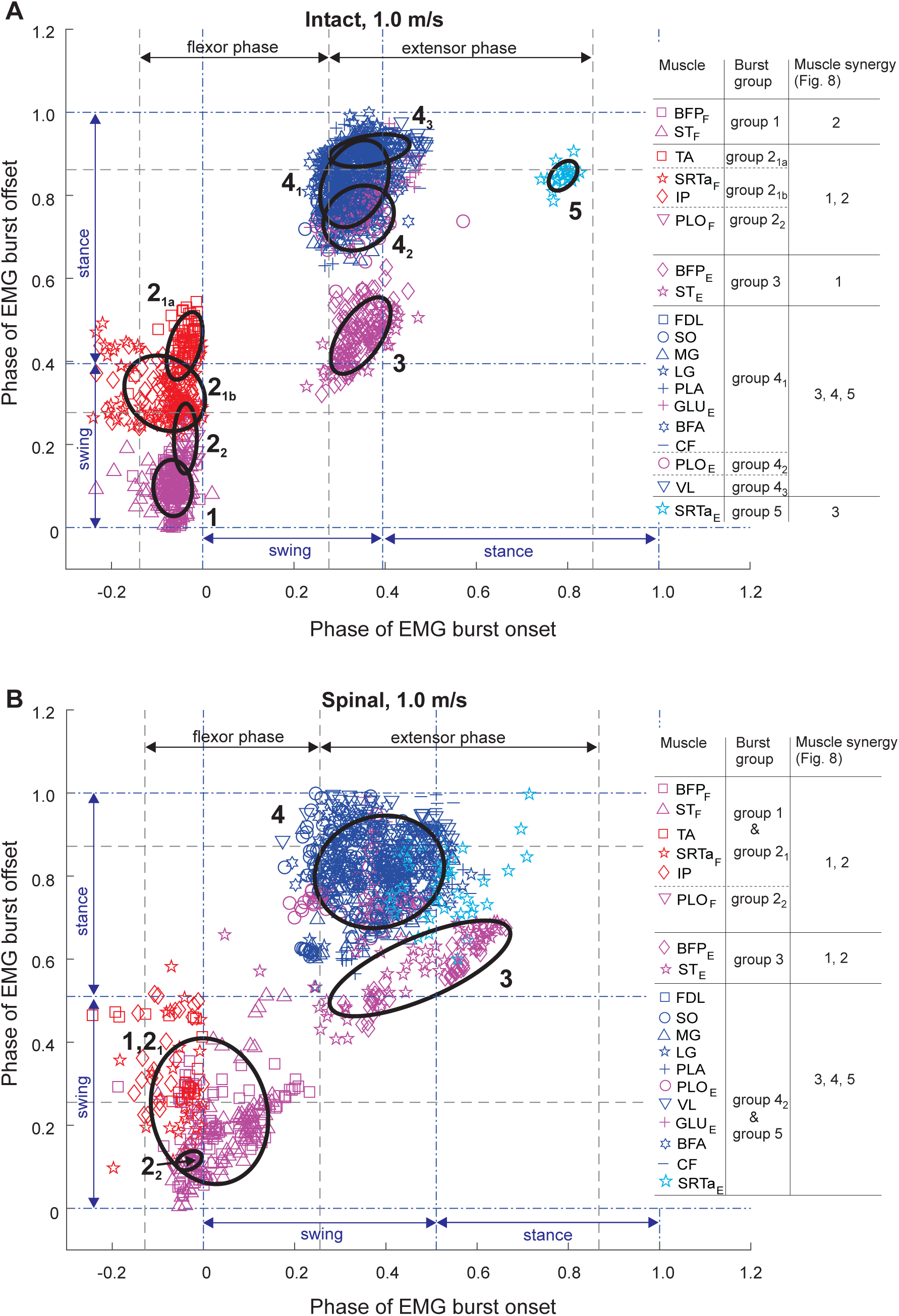
Scatter plots of EMG burst onset and offset times of 15 hindlimb muscles and EMG burst groups for locomotion speed of 1.0 m/s in intact (panel **A**) and spinal (panel **B**) states. For further information see the legend for Fig. 4.

### Muscle synergies and their activation patterns

*<u>Muscle synergies</u>*. EMG burst groups identified based on burst onset and offset times indicate muscles that are active at the same time, consistent with receiving a common neural input (Krouchev et al., 2006; Markin et al., 2012). Contributions of individual group members to the total EMG activity of the group and group activation patterns can be further assessed by a muscle muscle synergies that accounted for at least 90% of EMG variance in all muscles in each experimental condition. This number was found to be 5 in all experimental conditions (Fig. 7A). The main effect of state on the EMG variance accounted for (the coefficient determination R2) was insignificant (F_1,343_ = 2.6, p = 0.105), whereas the effects of speed and state-speed interaction were significant (F_2,343_ = 8.7, p < 0.001 and F_2,343_ = 15.6, p < 0.001, respectively). Although significant, the differences in R^2^ between the experimental conditions were rather small: the minimum and maximum values were 0.91 ± 0.02 (intact at 0.4 m/s) and 0.93 ± 0.02 (intact at 1.0 m/s). To evaluate the similarity of matrices **W** and **C** in different experimental conditions, we also computed the variance accounted for by different combinations of matrices 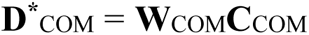 composed by mixing different experimental conditions (see Methods). As seen in Fig. 7B, 5 synergies were insufficient to obtain an R^2^ at or above 0.9 for any combination of experimental conditions. Seven synergies were necessary to obtain R^2^ > 0.9 for the combination intact-spinal condition at each speed, i.e.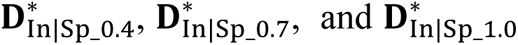 (first 3 bars for 7 synergies) and 8 synergies were required to exceed an R^2^ of 0.9 by the combination of all speeds for intact and spinal conditions, i. e. 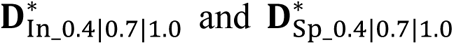. The combination of all 6 experimental conditions 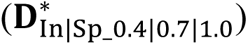 required 12 synergies to reach an R^2^ of 0.9 (Fig. 7B). These results suggest that 5 muscle synergies and/or their activation patterns found for each experimental condition differ between the conditions. Five muscle synergies (coefficients W) and their activation patterns (coefficients C) are shown in Fig. 8A **and B**, respectively, for each experimental condition and for the combination of all experimental conditions together. The order of muscle synergies was selected based on the appearance of the synergy’s activation peak in the locomotor cycle. Thus, the first two synergies had activity peaks at the onset and middle of swing, respectively, in all experimental conditions and in the combination of all conditions (Fig. 8B). The activation peak of synergy 3 occurred near the swing-stance transition, the activation peak of synergy 4 at early stance; and the activation peak of synergy 5 at middle-late stance in all experimental conditions. Muscle contributions to the corresponding synergies (coefficients W) are shown in Fig. 8A. Although the median of most coefficients W in all synergies and experimental conditions exceeded zero (nonparametric Wilcoxon signed-rank test, p < 0.001, n = 50), i.e. each muscle had a non-zero contribution to all synergies, we considered the weight coefficients W below 0.15 negligible (Klishko et al., 2021). The linear mixed-effect model analysis performed for individual synergies revealed significant effects on coefficients W of the factor muscle in all synergies (F_14,8820_ = 252.3-1161.4, p < 0.001), state in synergies 3 and 4 (F_14,8820_ = 252.3, p <0.001 and F_14,8820_ = 327.8, p < 0.001) but not synergies 1, 2, and 5 (F_14,8820_ = 1.8-2.4, p = 0.089-0.157), and speed in synergies 1, 3-5 (F_3,8820_ = 4.4-22.2, p ≤ 0.004) but not synergy 2 (F_3,8820_ = 1.5, p = 0.219).

**Figure 7.**
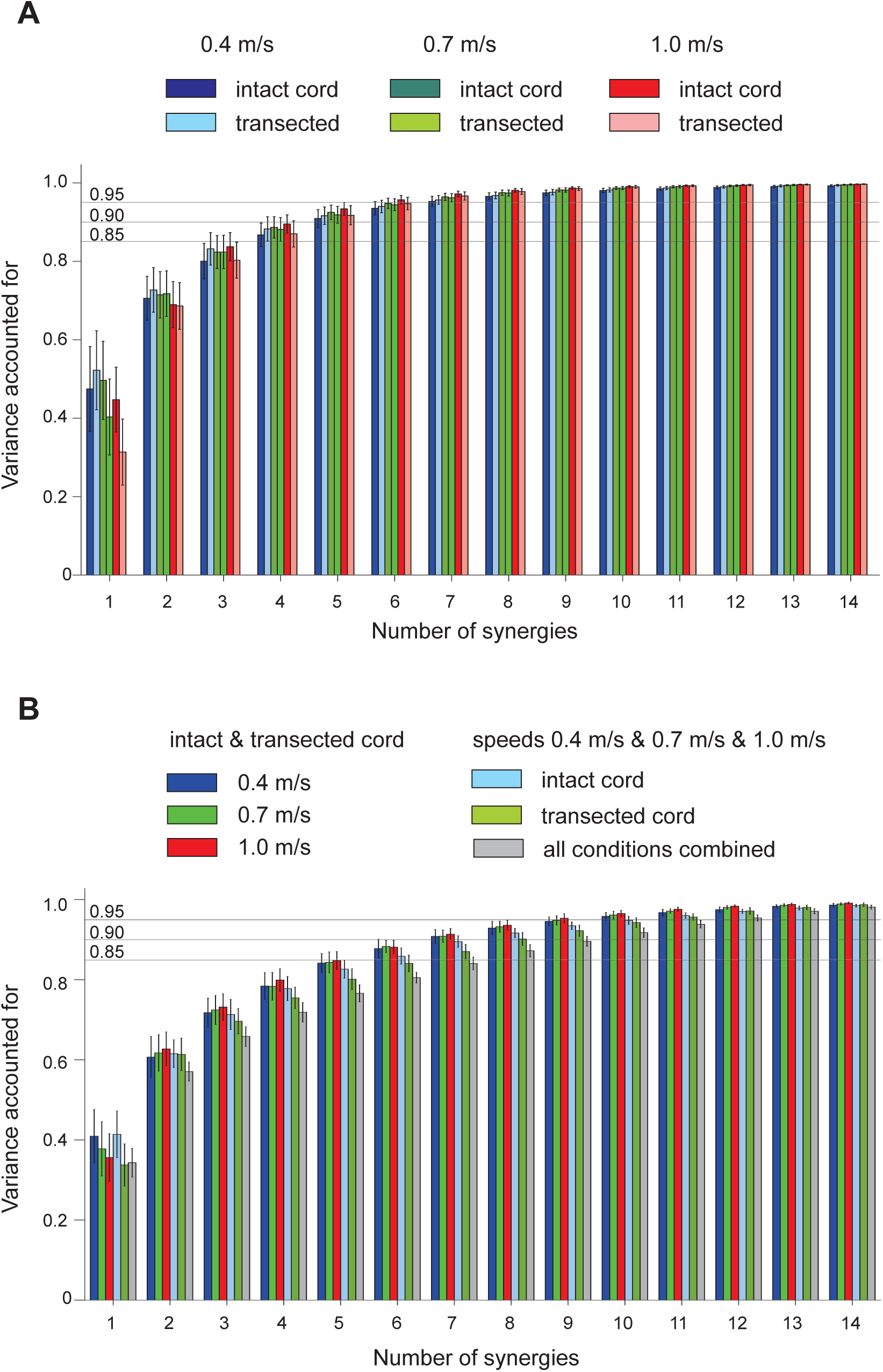
Variance accounted for (mean ± SD) in recorded EMG patterns by EMG patterns reconstructed from muscle synergies as a function of the number of synergies. **A**: Variance accounted for by reconstructed EMG patterns in each intact and spinal state during locomotion at 0.4, 0.7, and 1.0 m/s. Horizontal lines indicate values of variance 0.85, 0.90, and 0.95. **B**: Variance accounted for by reconstructed EMG patterns obtained from different combinations of matrices **D\***_COM_ = **W**_COM_**C**_COM_ composed by mixing different experimental conditions (see Methods). Variance computed for six combinations is shown: (1-3) the intact & spinal state combination at speeds 0.4, 0.7, and 1.0 m/s, respectively; (4-5) the combination of three speeds for intact and spinal state, respectively; (6) the combination of all 6 experimental conditions.

**Figure 8.**
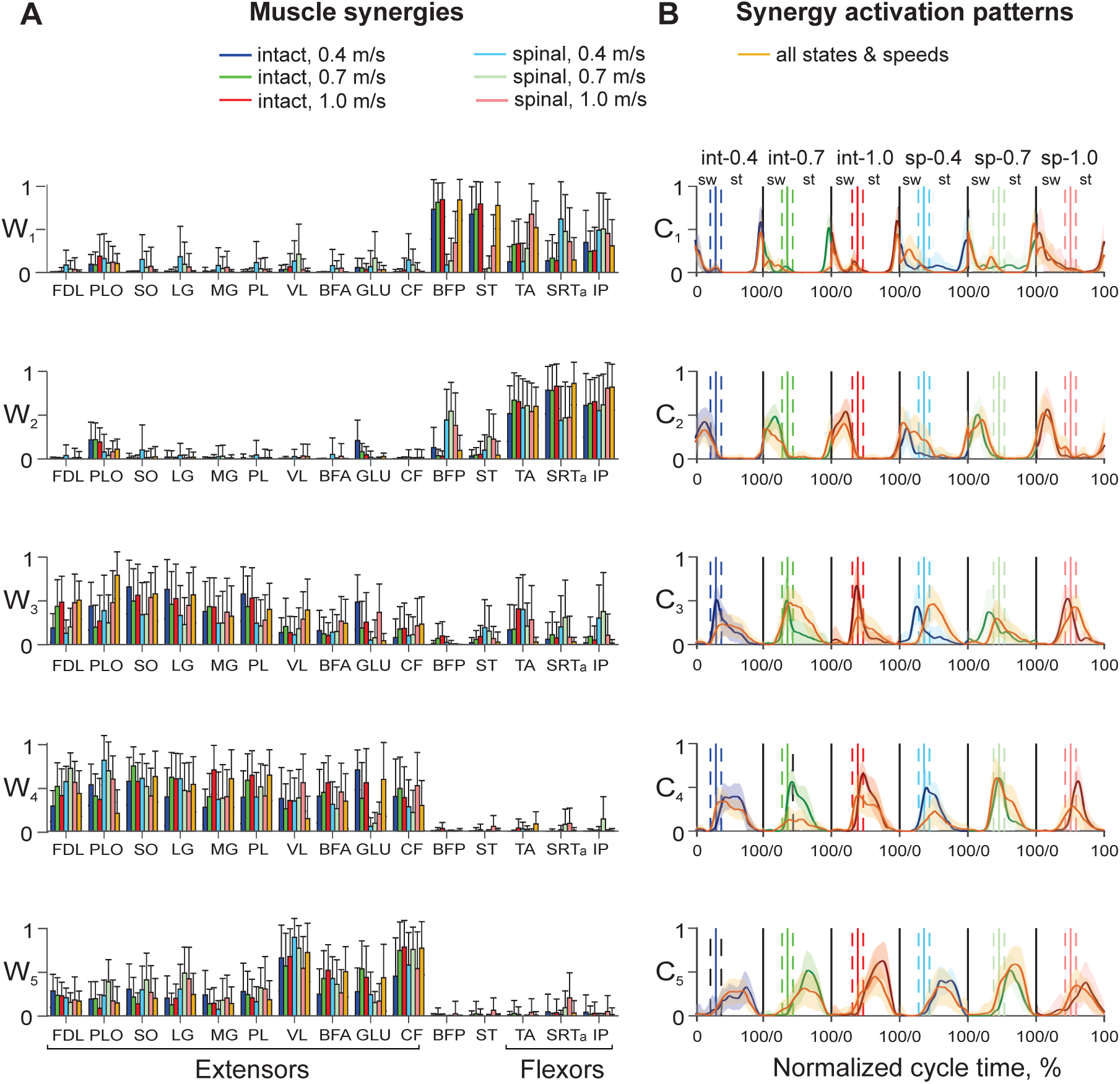
Five muscle synergies and their activation patterns during tied-belt locomotion at speeds 0.4, 0.7, and 1.0 m/s in intact and spinal conditions. **A**: Mean (±SD) muscle synergy weights (relative contribution of each muscle to a given synergy; rows in (*15* × *5*) matrices 𝐖_In_0.4_, 𝐖_In_0.7_, 𝐖_In_1.0_, 𝐖_Sp_0.4_, 𝐖_Sp_0.7_, 𝐖_Sp_1.0_ corresponding to each experimental condition: intact and spinal states at three locomotion speeds). Matrix 𝐖_In|Sp_0.4|0.7|1.0_ contains muscle weight coefficient for a combination of all 6 experimental conditions; matrix dimensions are (*15* × *5*). Each matrix **W** was computed using 50 randomly selected cycles across all cats (see Methods). **B**: Mean (±SD) activation patterns of five muscle synergies (columns in (*5* × *100*) matrices 𝐂_In_0.4_, 𝐂_In_0.7_, 𝐂_In_1.0_, 𝐂_Sp_0.4_, 𝐂_Sp_0.7_, 𝐂_Sp_1.0_ corresponding to each experimental condition. The activation patterns for each experimental condition are shown as a function of the normalized cycle time in the following order: intact (int), speed 0.4 m/s; intact, speed 0.7 m/s; intact, speed 1.0 m/s; spinal (Sp), speed 0.4 m/s; spinal, speed 0.7 m/s; and spinal, speed 1.0 m/s. Activation coefficients in matrix 𝐂_In|Sp_0.4|0.7|1.0_ (orange line) represent synergy activation patterns of a combination of all 6 experimental conditions; matrix dimensions are (*5* × *600*). Vertical continues lines surrounded by vertical dashed lines correspond to the mean ± SD swing offset/stance onset normalized times in each experimental condition. These lines separate the swing (sw) and stance (st) phases. For muscle abbreviations see the legend for Fig. 1.

Bifunctional muscles BFP and ST made the greatest contributions to synergy 1 during intact locomotion at all speeds (range 0.68 – 0.85) with much smaller contributions of flexors TA, SRTa, and IP (range 0.12 – 0.35). In spinal locomotion, these contributions were significantly smaller for BFP and ST (below 0.15 at speeds 0.4 and 0.7 m/s; 0.31 - 0.35 at 1.0 m/s), but larger for TA, SRTa, and IP at all speeds (0.28 - 0.68) with the exception of TA at speed 0.4 m/s (0.13). The contributions of extensor muscles or flexor bursts of bifunctional muscles to synergy 1 was negligible (below In synergy 2, flexors TA, SRTa, and IP significantly increased their contributions in the intact and spinal states at all three speeds (coefficients W ranged from 0.44 to 0.83, p ≤ 0.014) except for SRTa in spinal locomotion at speed 0.7 m/s (p = 0.868) and IP in spinal locomotion at speed 0.4 m/s (p = 0.186). The contributions of the bifunctional BFP and ST were negligible in intact locomotion at all speeds but significantly increased for BFP in spinal locomotion (range 0.38 – 0.45) compared to its contribution to synergy 1 at 0.4 and 0.7 m/s (p < 0.001) but not at 1.0 m/s (p = 0.421). The ST contribution to synergy 2 in spinal locomotion was small (0.10 – 0.25). The contribution of the PLO extensor burst was also small at all speeds in intact (range 0.19 – 0.22) and spinal (range 0.04 – 0.08) locomotion. The contribution of the extensor burst of GLU exceeded 0.15 only in intact locomotion at 0.4 m/s (0.21). The W coefficients of extensor muscles were below 0.15 and thus did not contribute substantially to synergy 2. Major contributors to synergies 1 and 2 (BFP, ST, TA, SRTa, and IP) corresponded to EMG burst groups 1, 2, and 3 (**Figs. 4-6**).

All extensor muscles contributed substantially to synergies 3, 4, and 5 during intact and spinal locomotion at all speeds. The greatest contributions to synergies 3 and 4 were provided by ankle extensors SO, MG, LG, and PL in intact (range 0.38 – 0.63) and spinal (0.21 – 0.54) locomotion (with significantly higher W coefficients in intact than spinal locomotion (p ≤ 0.05) with very number of exceptions. For example, the change was insignificant between the conditions for MG, synergy 3 at 1.0 m/s and synergy 4 at 0.7 m/s, p = 0.203 – 0.714; SO, synergy 4, speed 0.4 m/s, p = 0.458; and PL, synergy 4, speed 0.7 m/s, p = 0.105. The greatest contributions to extensor synergy 5 in all experimental conditions were provided by proximal muscles VL (range 0.542 – 0.896), BFA (0.248 – 0.519), GLU (0.152 – 0.535), and CF (0.452 – 0.785). Distal extensors contributed less to synergy 5; in intact locomotion their contribution was between 0.125 (LG, speed 0.7 m/s) and 0.304 (SO, speed 0.4 m/s). The contributions of flexors TA and IP and extensor bursts of SRTa, BFP, and ST to extensor synergies were substantial only in synergy 3, primarily in spinal locomotion, which ranged between 0.201 in SRTa at 0.4 m/s and 0.402 in TA at 0.4 m/s. The contributors to extensor synergies 3-5 corresponded to EMG burst groups 4 and 5 (**Figs. 4-6**).

We compared similarity of muscle contributions to each synergy between states and speeds by computing the scalar product (the angle) between pairs of 15-dimensional vectors of muscle weights (rows in **W**) for different combinations of experimental conditions (see Methods). The angle can theoretically change from 0° (the greatest similarity between the two vectors) to 90° (the greatest dissimilarity). The linear mixed-effects model analyses revealed significant effects of synergy (five levels), experimental conditions (2 states, 3 speeds, and combinations of states and speeds), and their interaction on the angles (F_4_,_3675_ = 150.8, p < 0.001; F_14_,_3675_ = 68.7, p < 0.001; and F_56_,_3675_ = 12.5, p < 0.001; respectively). The similarity of the muscle weight vectors for intact condition across speeds was modest for all synergies; angles ranged from 28.2 ± 10.7° to 40.5 ± 14.7° in synergies 1 and 2 and from 44.0 ± 18.5° to 58.4 ± 19.1° in synergies 3-5 (Fig. 9, first 3 bars in each synergy). The similarity of the muscle weight vectors between speeds in spinal locomotion was modest for synergies 2, 4, and 5 (38.7 ± 10.6° – 51.1 ± 14.6°) and relatively poor for synergies 1 and 3 (62.6 ± 13.7 – 67.9 ± 14.6) (Fig. 9, middle three bars in each synergy). Comparisons of the muscle weight vectors between intact and spinal locomotion at each speed revealed similar results, i.e. a modest similarity for synergies 2, 4, and 5 (43.3 ± 15.8° – 55.1 ± 18.3°) and relatively poor similarity for synergies 1 and 3 (56.5 ± 12.9° – 73.8 ± 15.8°) (Fig. 9, last When we compared muscle weight coefficients W between different combinations of states and speeds (various matrices **W**_COM_), we found generally similar distributions of muscle contributions to individual synergies compared to those revealed by synergy analysis of individual experimental conditions (compare Fig. 8A and Fig. 10). Specifically, flexor bursts of bifunctional BFP and ST had the greatest contributions to synergy 1 in the majority of state and speed combinations except for spinal locomotion at all speeds and the combination of intact and spinal locomotion at 0.7 m/s (Fig. 10). Flexor muscles TA, SRTa, and IP had generally lower contributions to synergy 1 but were main contributors to synergy 2 (typically above 0.5 and reaching a peak value of 0.922 ± 0.125 for SRTa in intact locomotion at all speeds). In extensor synergies 3-5, flexors and bifunctional BFP and ST had negligible contributions (in most cases below 0.15), whereas extensors and extensor bursts of PLO and GLU contributed substantially. As was the case in the individual experimental conditions (Fig. 8A), distal ankle extensors contributed more to synergies 3 and 4 (except for GLU whose contributions were between 0.359 and 0.837 for 4 out of 6 experimental combinations), while proximal extensors VL, BFA, GLU, and CF contributed to synergy 5 with weights typically exceeding 0.5 (Fig. 10).

**Figure 9.**
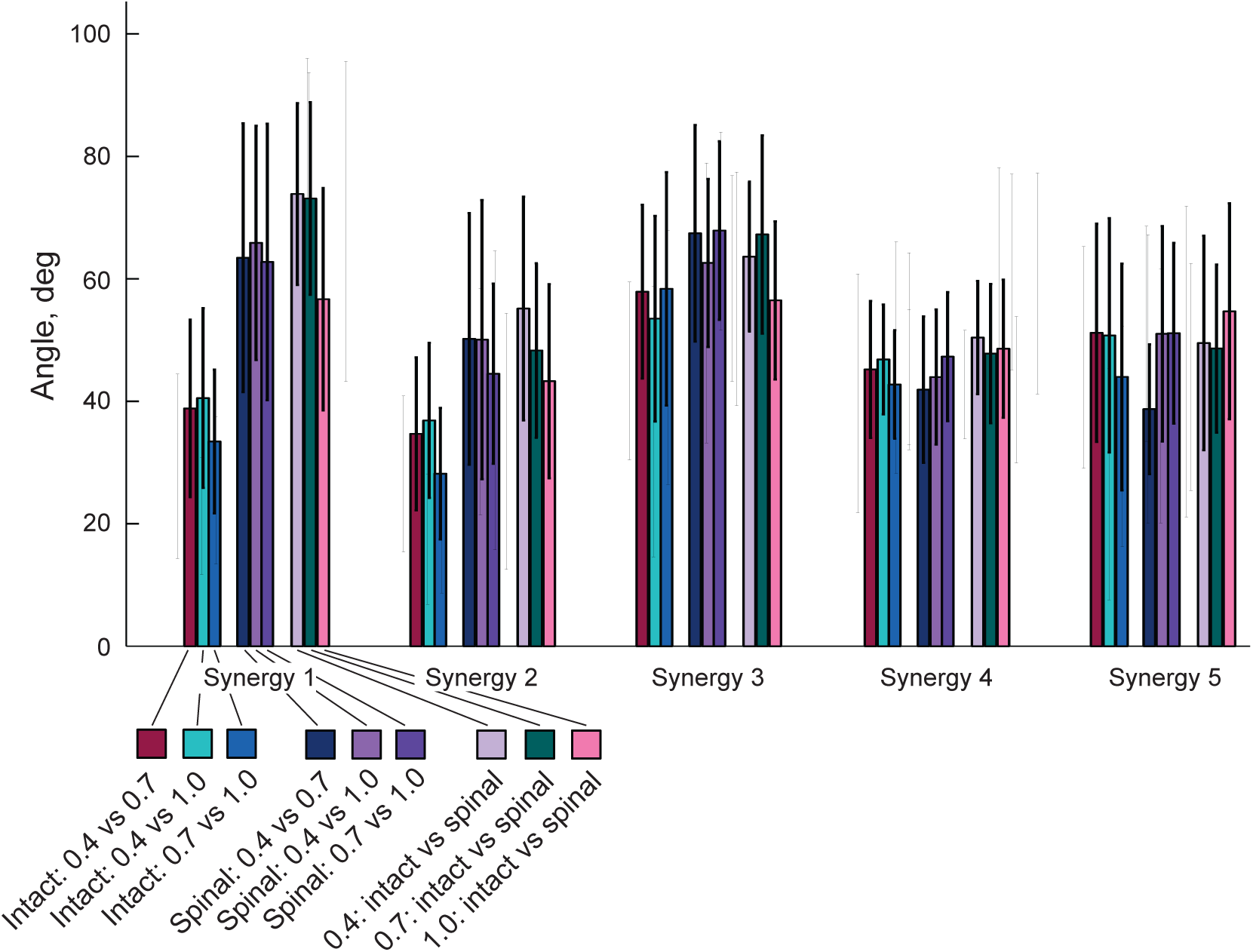
Angles (mean ± SD) between pairs of 15-dimensional vectors of muscle weights for a given synergy corresponding to all combinations of two different experimental conditions.

**Figure 10.**
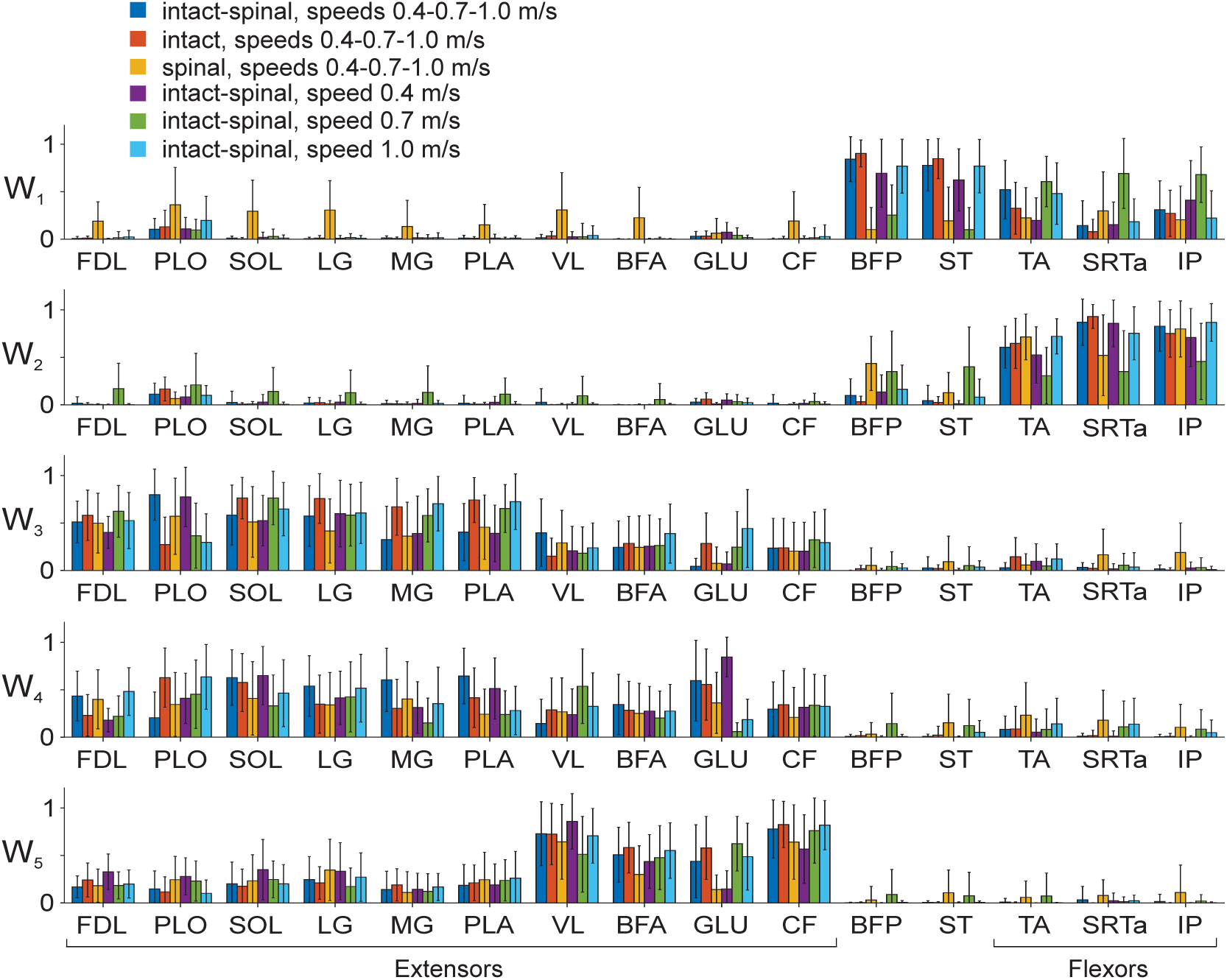
Five muscle synergies (mean ± SD) computed for different matrices **W**_COM_ composed by mixing different combinations of experimental conditions (see Methods). These combinations (intact, spinal, speeds 0.4, 0.7, and 1.0 m/s), matrix 𝐖_In|Sp_0.4|0.7|1.0_; (2) intact and the combination of all speeds, 𝐖_In_0.4|0.7|1.0_; (3) spinal and the combination of all speeds 𝐖_Sp_0.4|0.7|1.0_; (4) the combination of intact and spinal condition at speed 0.4 m/s, 𝐖_In|Sp_0.4_; (5) the combination of intact and spinal condition at speed 0.7 m/s, 𝐖_In|Sp_0.7_; and (6) the combination of intact and spinal condition at speed 1.0 m/s, 𝐖_In|Sp_1.0_. For muscle abbreviations see the legend for Fig. 1.

In summary, we found that 5 synergies were sufficient to account for at least 90% of the variance in EMG patterns in each of 6 experimental conditions. Although in all experimental conditions the first two synergies were flexor and the remaining 3 synergies, extensor, the composition (muscle weights) of the 5 synergies significantly differed among the intact and spinal states and locomotion speeds.

#### Synergy activation patterns

Activation patterns (time-dependent coefficients C, Fig. 8B) of all synergies were significantly affected by the factors time (F_9,11760_ = 709.2 – 2023.4, p < 0.001), state (F_3,11760_ = 29.2 – 97.6, p < 0.001), and speed (F_5,11760_ = 30.1 – 86.4, p < 0.001). In synergy 1, activation patterns of intact locomotion at different speeds were highly correlated, with mean coefficients of determination (R^2^) computed between all pairs of C coefficient vectors corresponding to different speeds between 0.68 and 0.69. These patterns were also highly correlated with the corresponding components of common matrix 𝐂_In_0.4|0.7|1.0_: R^2^ = 0.90-0.93 (Fig. 11A, orange lines, see Methods). The intact activation patterns for different speeds had a relatively low correlation with the corresponding spinal patterns (R^2^ = 0.30-0.49). In spinal locomotion, the activation patterns were dissimilar at different speeds as indicated by low R^2^ coefficients (0.27 – 0.28) and their correlation with the corresponding components of common matrix 𝐂_Sp_0.4|0.7|1.0_ was low: R^2^ = 0.21 – 0.37 (Fig. 11B). Components of common matrices 𝐂_In|Sp_0.4_, 𝐂_In|Sp_0.7_, 𝐂_In|Sp_1.0_, combining intact and spinal states at each speed, demonstrated high correlation only with the intact matrices 𝐂_In_0.4_ and 𝐂_In_1.0_ at 0.4 m/s (R^2^ = 0.90) and 1.0 m/s (R^2^ = 0.91) in synergy 1 (Fig. 11C, E, synergy 1). The spinal components of the above matrix **C** combining intact and spinal states demonstrated much lower correlations with the C coefficients for spinal locomotion at different speeds (R^2^ = 0.23-0.49, Fig. 11C**-E**).

**Figure 11.**
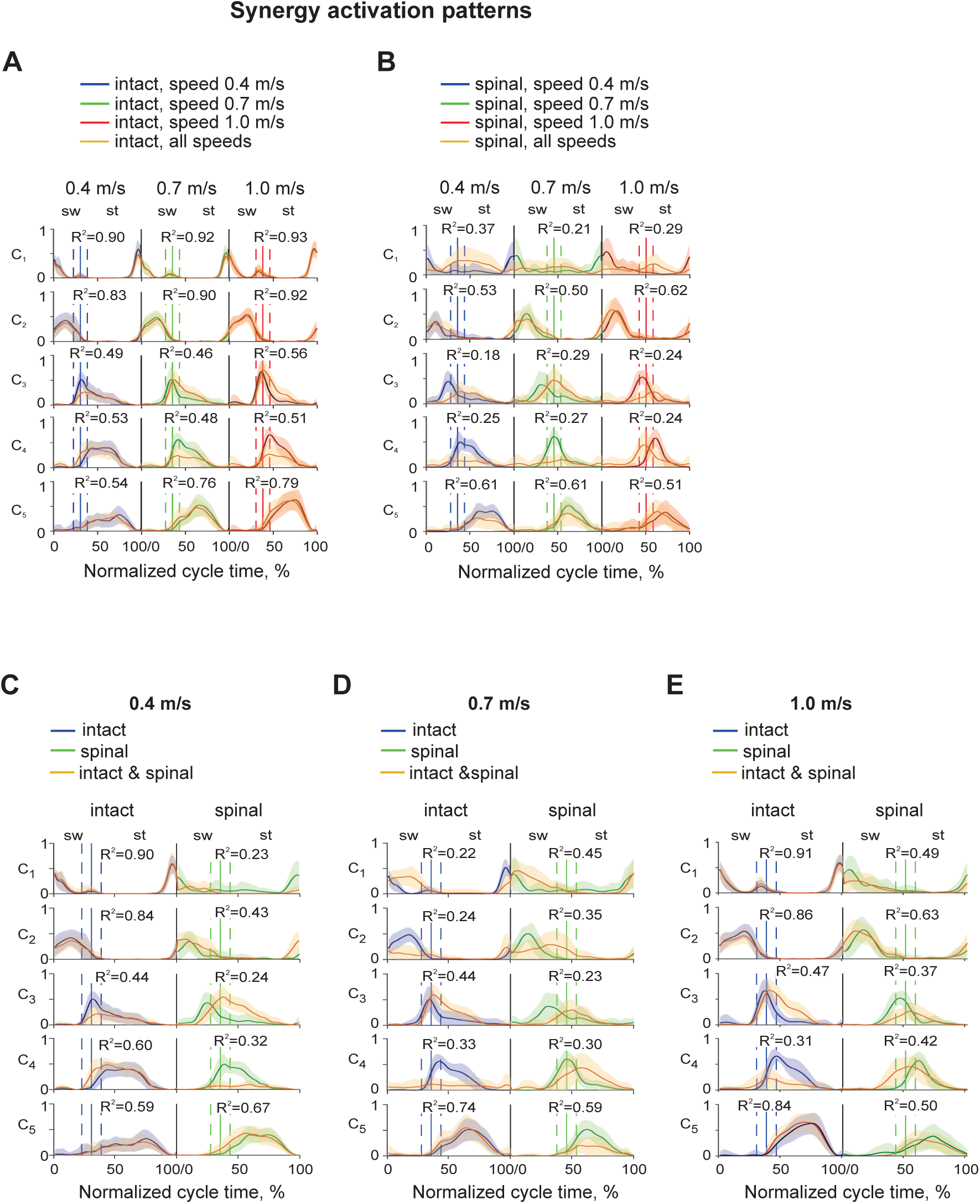
Synergy activation patterns (mean ± SD) as a function of the normalized cycle time for individual experimental conditions and their combinations. **A**: Activation patterns for intact state and different locomotion speeds are shown in the following order from left to right: speed 0.4 m/s, speed 0.7 m/s, and speed 1.0 m/s. Activation coefficients in the common matrix 𝐂_In_0.4|0.7|1.0_ (orange line) represent synergy activation patterns of a combination of all 3 speeds of intact state; the matrix dimensions are (*5* × *300*). Vertical continues lines surrounded by vertical dashed lines correspond to the mean ± SD swing offset/stance onset normalized times in each experimental condition. These lines separate the swing (sw) and stance (st) phases. Coefficients of determination (R^2^) indicate correlation between the activation pattern of an individual experimental condition with the corresponding component of the common matrix 𝐂_In_0.4|0.7|1.0_. **B**: Activation patterns for spinal condition and different locomotion speeds are shown in the following order from left to right: speed 0.4 m/s, speed 0.7 m/s, and speed 1.0 m/s. Activation coefficients in the common matrix 𝐂_Sp_0.4|0.7|1.0_ (orange line) represent synergy activation patterns of a combination of all 3 speeds of spinal condition. Coefficients of determination (R^2^) indicate correlation between the activation pattern of an individual experimental condition with the corresponding component of the common matrix 𝐂_Sp_0.4|0.7|1.0_. **C**: Activation patterns for intact and spinal states at locomotion speed of 0.4 m/s and the activation coefficients in the common matrix 𝐂_In|Sp_0.4_ (orange line). **D**: Activation patterns for intact and spinal states at locomotion speed of 0.7 m/s and the activation coefficients in the common matrix 𝐂_In|Sp_0.7_ (orange line). **D**: Activation patterns for intact and spinal states at locomotion speed of 1.0 m/s and the activation coefficients in the common matrix 𝐂_In|Sp_1.0_ (orange line).

Activation patterns of synergy 2 were generally consistent within and between states and speeds. Correlations for intact locomotion between speeds were R^2^ = 0.65-0.71 and R^2^ = 0.37-0.51 for spinal locomotion. Intact patterns at different speeds had a modest correlation with the 𝐂_In_0.7_, and 𝐂_In_1.0_ of intact locomotion (R^2^ = 0.83-0.92); similar correlations for spinal locomotion were lower (R2 = 0.50-0.62) (Fig. 11A, B). Correlations of components of common matrices 𝐂_In|Sp_0.4_, 𝐂_In|Sp_0.7_, 𝐂_In|Sp_1.0_ with the patterns for intact locomotion were medium-high at 0.4 and 1.0 m/s (R^2^ = 0.43-0.86) and low for speed 0.7 m/s (R^2^ = 0.24-0.35) (Fig. 11, C-E).

Activation patterns of extensor synergies 3-5 corresponding to different speeds were modestly correlated in intact locomotion (R^2^ = 0.43-0.66) and had a wide range of correlations in spinal locomotion from very low to modest (R^2^ = 0.08-0.63). Correlations between intact and spinal patterns at the same speeds were low to modest (R^2^ = 0.21-0.58). Correlations of components of common matrices 𝐂_In_0.4|0.7|1.0_ and 𝐂_Sp_0.4|0.7|1.0_ with the corresponding matrices C for individual states and speeds were much higher for intact locomotion (R^2^ = 0.46-0.79) than for spinal locomotion (R^2^ = 0.18-0.61) (Fig. 11A, B). Correlations of components of common matrices 𝐂_In|Sp_0.4_, 𝐂_In|Sp_0.7_, 𝐂_In|Sp_1.0_ with the corresponding individual matrices **C** were modest-high for intact locomotion (R^2^ = 0.44-0.84) and low-modest for spinal locomotion (R^2^ = 0.23-0.67) at all speeds (Fig. 11C**-E**).

In summary, activation patterns of each synergy in intact locomotion at different speeds were similar to each other and to the components of the common matrices **C**_COM_ with very few exceptions. Activation patterns of spinal locomotion, on the other hand, were mostly different across speeds and dissimilar compared to the components of the common matrices **C**_COM_. Nevertheless, the phases of high activation and the appearance of the activation peaks in the cycle were consistent across states and speeds (Figs. 8B, 1**1**).

## Discussion

The goal of this study was to determine hindlimb muscle synergies and their activation patterns during walking at different locomotion speeds before and after spinal transection. The obtained results supported our hypothesis that a spinal mechanism controls the number of muscle synergies as a function of speed during locomotion. We found that 5 synergies were sufficient to account for 90% of the EMG variance in each of 6 experimental conditions (i.e. 2 states x 3 speeds). However, the obtained muscle contributions to each synergy (weights W) and the synergy activation patterns (time-dependent coefficients C) in the different experimental conditions did not support our hypothesis that a spinal mechanism fully explains the composition and activation patterns of muscle synergies during locomotion at different speeds.

### State-dependent effects on muscle synergies

Previous studies of EMG burst groups and muscle synergies during tied-belt treadmill locomotion after low-thoracic spinal transection demonstrated that hindlimb muscle synergies in adult cats originate in the lumbar spinal cord (Desrochers et al., 2019; Higgin et al., 2020; Harnie et al., 2021) and are generally similar to the EMG burst groups and synergies during intact cat locomotion (Krouchev et al., 2006; Markin et al., 2012; Klishko et al., 2021). The results of the present study are generally consistent with previous conclusions. We also found between 5 and 9 In the present study, spinal transection reduced the number of subgroups from 7 to 5 at a speed of 0.4 m/s by merging group 1 and subgroup 2_1_ (Fig. 4), from 7 to 6 at 0.7 m/s by merging subgroup 4_2_ and group 5 (Fig. 5), and from 9 to 4 at 1.0 m/s by merging group 1 with subgroup 2_1_ and subgroup 4_2_ with group 5 (Fig. 6). These results are consistent with a previous study (Higgin et al., 2020) reporting a decrease in the number of burst groups from 5 to 2 during level intact and spinal locomotion at 0.4 m/s, respectively. Another study (Desrochers et al., 2019) also demonstrated a more compact grouping of muscle bursts that included SRTa and tensor fascia lata after spinal transection due to changes in burst onsets/offsets in these muscles.

We also found a substantial shift in the normalized EMG onset time in extensors around mid-swing (Fig. 1). This shift in the flexor-extensor phase transition after spinal transection could result from the state-machine operation regime of spinal networks that depends on motion-dependent sensory feedback from the hindlimbs for phase transitions (Rybak et al., 2024). Indeed, previous studies of hindlimb-only treadmill locomotion in spinal cats demonstrated a maximum hip flexion much earlier in the swing phase, closer to mid-swing, than intact cats [see Fig. 3 in (Harnie et al., 2022)]. Maximum hip flexion corresponds to the maximum length of the hamstring muscles (Gregor et al., 2006; Klishko et al., 2021) and peak activity of group I and II muscle spindle afferents from these muscles (Prochazka and Gorassini, 1998). The occurrence of peak hip flexion and hamstring length in the swing phase is strongly correlated with the onset of hindlimb extensor activity (McVea et al., 2005; Gregor et al., 2006).

The general composition of the muscle synergies (synergies 1 and 2 and 3-5 for flexors and extensors, respectively) and their activation patterns were qualitatively similar in intact and spinal locomotion (Figs. 8, 10, 11). However, more in-depth analyses (statistical tests, angles between muscle weight coefficient vectors, and matrices **W**_COM_ and **C**_COM_) revealed significant differences in both synergy composition and activation patterns (**Figs. 9-11**). These differences are likely caused in part by changes in spinal neuron properties (membrane receptor and neurotransmitter production, synaptic connections, dendritic sprouting, etc.) and in afferent and propriospinal pathways due to spinal plasticity after spinal transection (Murray et al., 2010; Rossignol and Frigon, 2011; Bilchak et al., 2021; Martin, 2022; Punjani et al., 2023). Changes in hindlimb locomotor mechanics and thus in motion-dependent somatosensory feedback also undoubtedly affect the composition and/or activation patterns of muscle synergies (Santuz et al., 2019).

Muscle synergies and their activation patterns during spinal locomotion in this study are much more consistent with intact condition than typically reported in studies of people with spinal cord injury (Fox et al., 2013; Hayes et al., 2014; Danner et al., 2015; Sun et al., 2022). This is likely related to difficulties of recruiting patients with the same spinal location and severity of spinal injury, rehabilitation interventions, general health status, etc. It is also easier to control the posture of spinal cats when they step on the treadmill. Nonetheless, major features of muscle synergies in spinal cats and people with spinal cord injury are common and include flexor and extensor synergies, and the inclusion of individual synergies with muscles acting across several joints. The common features of muscle synergies suggest a similar organization of spinal locomotor networks in cats and humans.

#### Speed-dependent effects on muscle synergies

Locomotor speed did not affect general features of muscle EMG burst groups (**Figs. 4-6**) and muscle synergies (number, general composition, and pattern, Figs. 7A, 8). However, increasing speed led to an increase in the normalized magnitude of extensor and flexor EMG bursts (Fig. 2), changes in the composition of muscle burst subgroups within generally the same burst groups (**Figs. 4-6**), changes in muscle synergy weights (Fig. 8A), and shifting phases and increasing peak magnitude of synergy activation patterns (Fig. 8B) in both intact and spinal locomotion. All these changes are expected given an increased supraspinal drive and motion-dependent somatosensory feedback in intact locomotion and somatosensory feedback in spinal locomotion with increasing speed (Shik et al., 1966; Forssberg et al., 1980; Frigon et al., 2017; Caggiano et al., 2018).

Our results are consistent with human studies that demonstrated small variability in the number and composition of muscle synergies, with greater changes in synergy activation patterns within the same gait, walking or running (Ivanenko et al., 2003; Monaco et al., 2010; Gui and Zhang, 2016; Yokoyama et al., 2016; Saito et al., 2018; Dewolf et al., 2019). Substantial differences in muscle synergy composition and/or their activation patterns have been reported between walking and running in humans (Ivanenko et al., 2008; Hagio et al., 2015; Yokoyama et al., 2016). Five out of 9 cats in spinal locomotion switched to a running gait at speed 1.0 m/s in our study. Although we did not notice abrupt changes in muscle synergies when this occurred, we did find more flexor EMG bursts with greater magnitude than in intact locomotion at 1.0 m/s, where a walking gait was maintained (Fig. 2B). This result is consistent with previous observations that increased demands on leg flexors during the swing phase trigger the walk-run transition (Prilutsky and Gregor, 2001; Hreljac et al., 2007; Ivanenko et al., 2008).

### Potential organization of the spinal locomotor network controlling a single hindlimb

The number and composition of muscle burst groups and synergies obtained in this study provide information on the possible organization of the spinal locomotor network controlling hindlimb muscle activity. Two flexor and three extensor synergies that include muscles spanning ankle, knee and hip joints appear consistent with a two-level CPG organization proposed previously (Rybak et al., 2006a; McCrea and Rybak, 2008). The pattern formation level (**PF**) of such a model can contain 5 premotor interneuronal populations forming 5 muscle synergies. Two of these populations receive excitatory inputs from the flexor rhythm generator (**RG**) half-center and motion-dependent somatosensory feedback, while the other 3 PF populations receive excitatory inputs from the extensor RG half-center and motion-dependent somatosensory feedback (Fig. 12). According to the two-level CPG organization, the somatosensory feedback received by PF interneuronal populations modulates the duration and magnitude of activity in flexor and extensor motoneurons without changing or resetting the locomotor cycle duration (non-resetting feedback). Changes in locomotor rhythm can be accomplished by somatosensory feedback received by the RG half-centers (resetting feedback, Fig. 12). The reciprocal inhibition between the PF flexor and extensor populations reflects a lack of coactivation between the activation patterns of flexor synergies 1 and 2 and extensor synergies 3-5 (Fig. 8B). Reflex pathways with Ia inhibitory interneurons (not shown) provide additional contribution to reciprocal activity of flexors and extensors (Feldman and Orlovsky, 1975; Geertsen et al., 2011). The PF flexor and extensor muscles with different weights. Motoneurons controlling bifunctional muscles with both flexion and extension actions at different joints receive excitatory inputs from both the PF flexor and extensor populations as suggested previously (Perret and Cabelguen, 1980; Prilutsky, 2000; Shevtsova et al., 2016). The preliminary computer simulations using our neuromechanical model of cat hindlimb spinal locomotion utilizing a similar organization of the CPG and somatosensory afferent inputs (Markin et al., 2016; Rahmati et al., 2023) reproduced basic kinematic and muscle activity patterns recorded experimentally.

**Figure 12.**
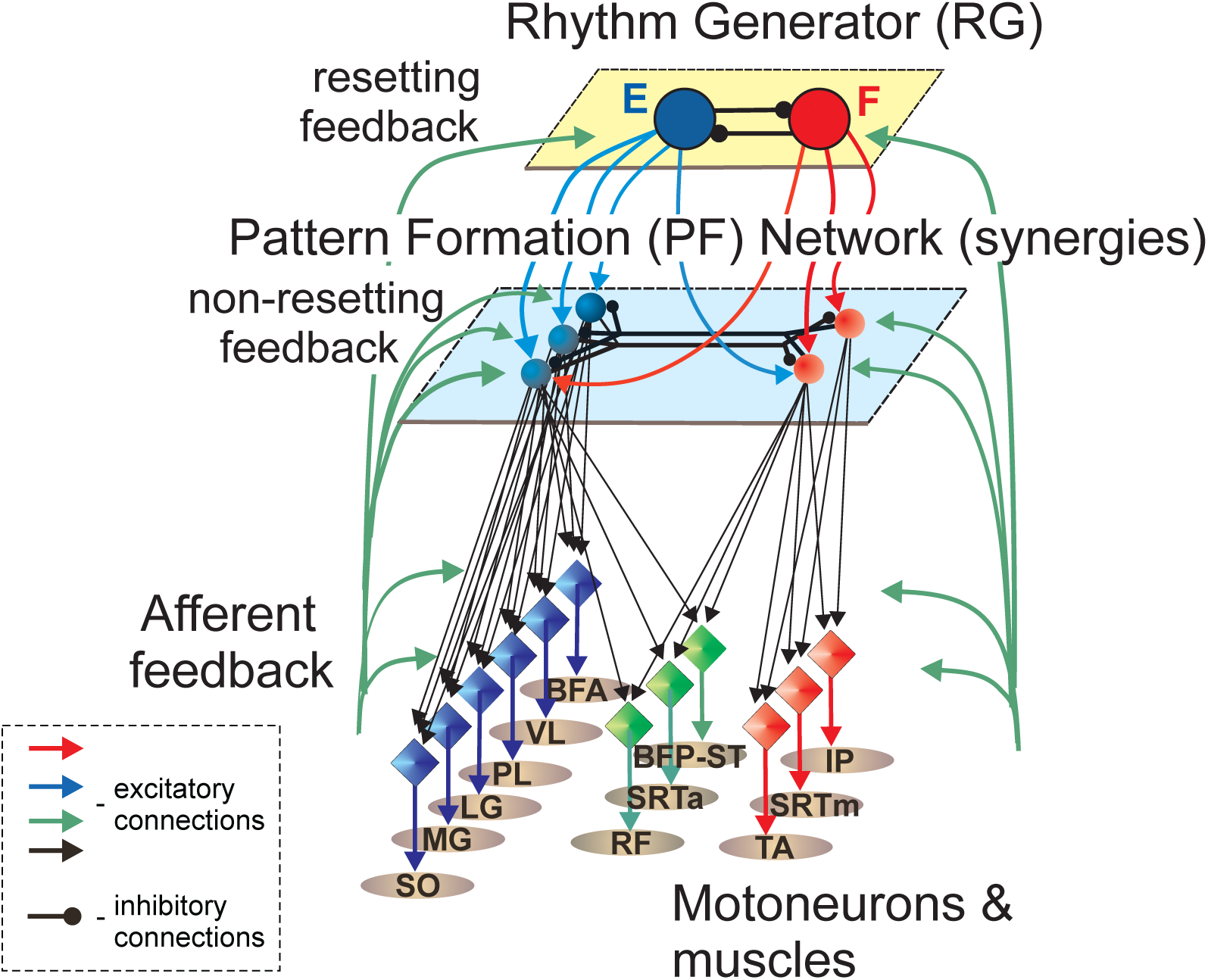
Schematic of a spinal locomotor CPG for a single hindlimb consistent with five synergies revealed in spinal condition. The schematic is a modified version of a previous two-level CPG model (Rybak et al., 2006b; McCrea and Rybak, 2008). The model consists of a rhythm generator with flexor and extensor half-centers (large circles) mutually inhibiting each other, and a pattern formation network that contains 5 premotor interneuronal populations (smaller circles) activating motoneurons (diamonds) of the corresponding muscles (ovals). For muscle abbreviations see the legend for Fig. 1. Additional muscle legends: RF, rectus femoris (hip flexor, knee extensor) and SRTm, sartorius medial (hip flexor and knee flexor). These muscles were not investigated in this study and included in the schematic based on previous synergy analyses (Markin et al., 2012; Higgin et al., 2020; Klishko et al., 2021). See text for further explanations.

## Conclusions

To better understand the organization of spinal locomotor networks, we investigated the effects of low thoracic spinal cord transection and locomotion speed on EMG activity, EMG burst groups, and muscle synergies in cats during treadmill locomotion at different speeds. We found that both the spinal transection and locomotion speed changed the durations of cycle and phase durations, as well as the EMG burst magnitude of hindlimb flexors and extensors, and the normalized EMG burst onset and offset times. The major EMG burst groups, the number of muscle synergies, and major features of muscle synergy composition and activation patterns were not substantially affected by spinal transection and locomotion speed, consistent with a spinal control mechanism. In all experimental conditions, we typically observed (i) 5 major EMG burst groups of hindlimb flexors and extensors; (ii) two muscle synergies composed of BFP and/or ST bifunctional muscles and flexors active at the stance-swing transition and during the swing phase, respectively; and (iii) three extensor synergies composed of hindlimb extensors and active at the swing-stance transition, the early stance phase, and the mid-late stance phase, respectively. Both spinal transection and locomotion speed modified subgroups of the EMG burst groups and the composition and activation patterns of selected synergies. The obtained results suggest that supraspinal drive, plastic changes in the spinal cord after its transection, and motion-dependent sensory feedback do not modify the general organization of the spinal locomotor networks but rather produce more subtle changes in, for example, neuronal properties, synaptic connections, somatosensory and propriospinal pathways. Based on the obtained results, we proposed an organization of a PF network of a two- level CPG, which can be tested in neuromechanical simulations of cat spinal locomotion.

## Acknowledgements

The authors thank Ateendra Subramanian and Ashley V. Nguyen for their help with data analysis.

## Author contributions

A.F., B.I.P., and I.A.R. conceived and designed research. J.H. and A.F. performed surgeries and data collection. A.N.K. developed computer code. A.N.K., C.E.H., and S.M.R performed data analysis. A.N.K. and B.I.P. prepared figures. B.I.P. drafted manuscript. All authors discussed, reviewed, edited, and approved the final version of the manuscript.

